# Setting the SCENE for Interpretable Cell–Gene Embeddings in Single-Cell RNA-seq

**DOI:** 10.64898/2026.09.12.750699

**Authors:** Oscar Lauritz Møberg, Magnus Blom Petersen, Tue Herlau, Lars Erik Kristensen, Leon Eyrich Jessen, Morten Mørup

**Author notes:** These authors jointly supervised this work and share senior authorship (Leon Eyrich Jessen, Morten Mørup).

## Abstract

Single-cell RNA sequencing measures cellular states at high resolution, but sparse high-dimensional count data remain difficult to model interpretably. We introduce the Single-Cell Euclidean Network Embedding (SCENE), a probabilistic latent-distance model that jointly embeds cells and genes from Unique Molecular Identifier (UMI) counts. SCENE treats the count matrix as a weighted bipartite cell-gene graph, where Euclidean distances represent transcriptional affinity, and combines this geometry with a zero-inflated count likelihood that separates gene detection from expression magnitude. Across real and simulated scRNA-seq datasets, SCENE recovers biologically structured cell and gene embeddings with state-of-the-art performance. Surprisingly, major biological structure is preserved in native two- and three-dimensional latent spaces, enabling directly interpretable visualization. Perturbation analyses show that SCENE organizes glucocorticoid-response genes and T-cell receptor regulatory programs coherently in gene space, capturing biology beyond cell-type separation. SCENE provides a transparent representation learning framework in which low-dimensional Euclidean geometry supports accurate modeling and biological interpretation.

## Introduction

Single-cell RNA sequencing (scRNA-seq) is a valuable and rapidly expanding methodology with wide use across health science and biology. By resolving gene expression at cellular resolution, scRNA-seq provides high-resolution measurements of cellular states and has made it possible to characterize cellular heterogeneity, identify rare subpopulations, and map dynamic transcriptional states, providing insight into both basic biology and complex disease mechanisms [1, 2]. At the same time, scRNA-seq data are high-dimensional, sparse, heterogeneous, and affected by substantial technical variability, creating persistent challenges for computational analysis. Current computational frameworks therefore often struggle to fully exploit sparse count data while retaining interpretability [3, 4].

Most single-cell workflows begin by normalizing UMI counts, applying a log transformation, selecting highly variable genes, and reducing dimensionality before downstream analysis [5, 6]. These steps are useful in practice, but they also impose modeling assumptions before the main representation is learned. In particular, normalization and feature selection can alter mean–variance relationships, attenuate biologically meaningful variation, and discard information that may be relevant for downstream inference [7, 6, 8]. Because UMI are discrete molecular counts, a natural alternative is to model the count-generating process directly. Count-aware likelihoods can combine variance modeling, noise correction, and signal extraction within a single statistical framework, reducing reliance on irreversible preprocessing transformations [5, 9, 10, 11].

A second challenge is representational. Many methods reduce scRNA-seq data to cell-level embeddings or cell– cell graphs constructed after preprocessing, often from PCA or other low-dimensional feature representations. Although effective, these representations can inherit biases from the preprocessing pipeline and may obscure the gene-level features that define cellular relationships [12, 13, 14]. Graph-based and encoder-based models have expanded the range of available representations, including methods that use neural encoders, graph neural networks, or bipartite cell–feature structure [15, 16, 17, 18, 19]. However, these approaches often trade off raw-count modeling, explicit cell-gene relational structure, and direct interpretability of the learned parameterization [20, 21].

These considerations suggest several desiderata for scRNA-seq representation learning. A model should preserve the count nature of UMI data and place the mean– variance relationship in an explicit likelihood where possible [7, 6, 5, 22]. It should minimize irreversible preprocessing assumptions or make their statistical consequences explicit [7, 6, 8, 14]. It should retain interpretable relationships between cells and molecular features rather than reducing the problem exclusively to a cell–cell graph [15, 18, 19]. Due to the application of scRNA-seq for discovery, the model structure should also support interpretation of the learned representation [20, 21]. Finally, because comparative single-cell studies are strongly affected by technical variability, the model should explicitly account for observed technical covariates while preserving biological variation [10, 23, 24, 17]. Table 1 summarizes how representative methods instantiate these desiderata.

**Table 1.** Comparison of representative scRNA-seq methods. Methods are compared by whether they provide a directly interpretable parameterization, learn a joint embedding of cells and features, operate directly on raw count data without prior normalization or log-transformation, avoid prior data reduction, use an explicit count likelihood, define an entry-wise generative model for each cell–gene observation, and explicitly account for batch effects or observed covariates. ✓, supported; ✗, not supported; *◦*, partially supported or task dependent.

| Method | Directly interp. | Joint embed. | Raw counts | No data reduction | Count likelihood | Generative model | Batch aware |
| --- | --- | --- | --- | --- | --- | --- | --- |
| SIMBA [18] | ○ | ✓ | ✗ | ✓ | ✗ | ✗ | ✓ |
| scVI [17] | ✗ | ✗ | ✓ | ✓ | ✓ | ✓ | ✓ |
| PCA/Harmony [23] | ○ | ✗ | ✗ | ✗ | ✗ | ✗ | ✓ |
| scBiG [19] | ✗ | ✓ | ✗ | ✓ | ✓ | ✓ | ✗ |
| scLDM [27] | ✗ | ✗ | ✓ | ✓ | ✓ | ✓ | ✓ |
| SCENE (ours) | ✓ | ✓ | ✓ | ✓ | ✓ | ✓ | ✓ |

To address these desiderata, we introduce the Single-Cell Euclidean Network Embedding (SCENE), a probabilistic latent-distance model that jointly embeds cells and genes directly from raw scRNA-seq counts. SCENE is built on the view that the count matrix can be treated as a bipartite relational system between cells and genes, where cells and genes occupy a shared Euclidean latent space and each observed cell–gene count is modeled as a function of the distance between their latent positions [25]. SCENE combines this latent-distance geometry with a zero-inflated Poisson likelihood and cell- and gene-level random effects [26], enabling count-aware representation learning, which gives the cell–gene distances a direct interpretation as transcriptional affinity, i.e. the distance-based relational importance of a gene to a cell beyond overall cell depth and gene abundance, while also allowing for multi-layer batch correction. Across diverse scRNA-seq datasets, we show that SCENE improves biological conservation and integration performance while producing stable, interpretable embeddings that remain biologically meaningful even in native ultra-low-dimensional (i.e., two- and three-dimensional) spaces. Importantly, this geometry also extends beyond cell identity as perturbation analyses show that SCENE recovers coordinated glucocorticoid-response modules and resolves T-cell receptor (TCR)-associated transcriptional, cytokine-response, and metabolic regulatory programs more clearly than SIMBA [18], a competing joint embedding approach.

These results highlight that SCENE is not only an integration and representation-learning framework, but also a tool for interpreting dynamic cell–gene and gene–gene relationships directly within the geometry of the native representation.

## Results

### The SCENE Model

SCENE can be viewed as a direct geometric model of the observed UMI count matrix itself. Each entry in this matrix records the number of UMIs observed for a given gene in a given cell (Fig. 1a). Rather than normalizing, transforming, and reducing the data to a cell-only representation, SCENE treats the count matrix as a weighted bipartite graph, with cells and genes as separate node classes and observed counts defining the cell-gene edge weights (Fig. 1b). The model embeds both cells and genes into a shared Euclidean latent space, where shorter cell–gene distances correspond to stronger transcriptional affinity, reflected in higher detection probabilities and larger expected counts, while cell- and gene-specific effects account for baseline variation such as sequencing depth and gene abundance (Fig. 1c). This makes the learned representation both a statistical model of the counts and an interpretable map of cell–cell, cell–gene, and gene–gene relationships. We use this shared geometry to evaluate whether SCENE recovers known cell structure, preserves perturbation responses, integrates across batches, remains interpretable in native low-dimensional spaces, and organizes biologically coherent gene programs across seven publicly available biological datasets and two simulated datasets (see Methods).

**Figure 1.**
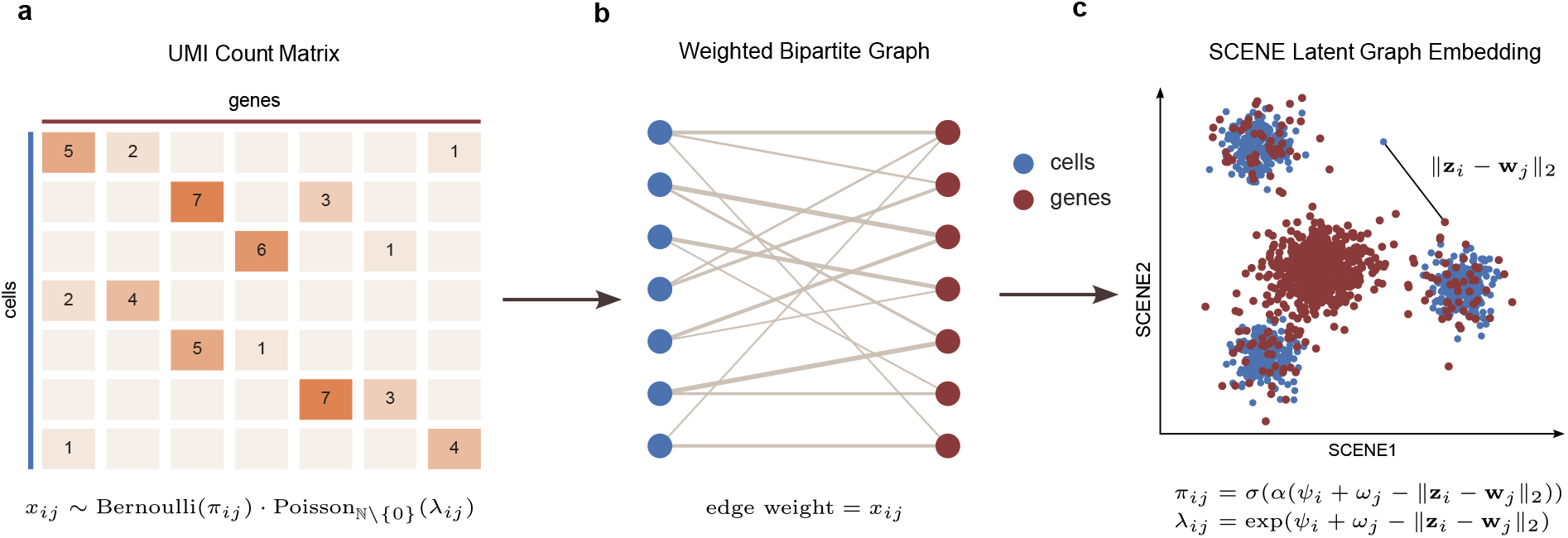
SCENE models raw UMI counts as a weighted cell-gene graph and embeds cells and genes in a shared Euclidean latent space. **a**, Raw UMI count matrix, where each entry *x*_*ij*_ represents the observed count for gene *j* in cell *i* and is modeled with a zero-inflated count likelihood. **b**, The count matrix is represented as a weighted bipartite graph with cells and genes as disjoint node classes, and edge weights given by the observed counts *x*_*ij*_. **c**, SCENE learns joint latent positions for cells and genes in a Euclidean metric space, where cell-gene distance directly determines both the probability of nonzero expression and the expected expression magnitude, while cell- and gene-specific random effects capture baseline variation. Here, **z**_*i*_ and **w**_*j*_ denote cell and gene positions; *Ψ*_*i*_ and *ω*_*j*_ are cell- and gene-specific effects; *λ*_*ij*_ is the underlying Poisson rate; *π*_*ij*_ is the probability of a nonzero count; *α* is a learned scaling parameter; and *σ* denotes the sigmoid function.

### SCENE learns a biologically structured joint cell-gene latent space

To assess whether SCENE learns a biologically meaningful joint latent representation, we examined the correspondence between unsupervised clustering and reference cell-type annotations in the PBMC CITE-seq dataset [12]. We applied agglomerative clustering with Ward linkage to the learned joint latent space and selected 18 clusters, corresponding approximately to log_2_(*N* + *G*), where *N* and *G* denote the numbers of cells and genes, respectively (Fig. 2a,b). Comparison with the reference annotations shows strong agreement across major cell types, with most assignment discrepancies occurring within closely related T-cell and monocyte-associated populations rather than between unrelated lineages (Fig. 2d). Moreover, the use of Laplacian initialization (see Methods) already recovers broad biological structure, which is largely preserved after optimization and remains consistent with both the final cluster assignments and annotated cell types (Fig. 2b,c).

**Figure 2.**
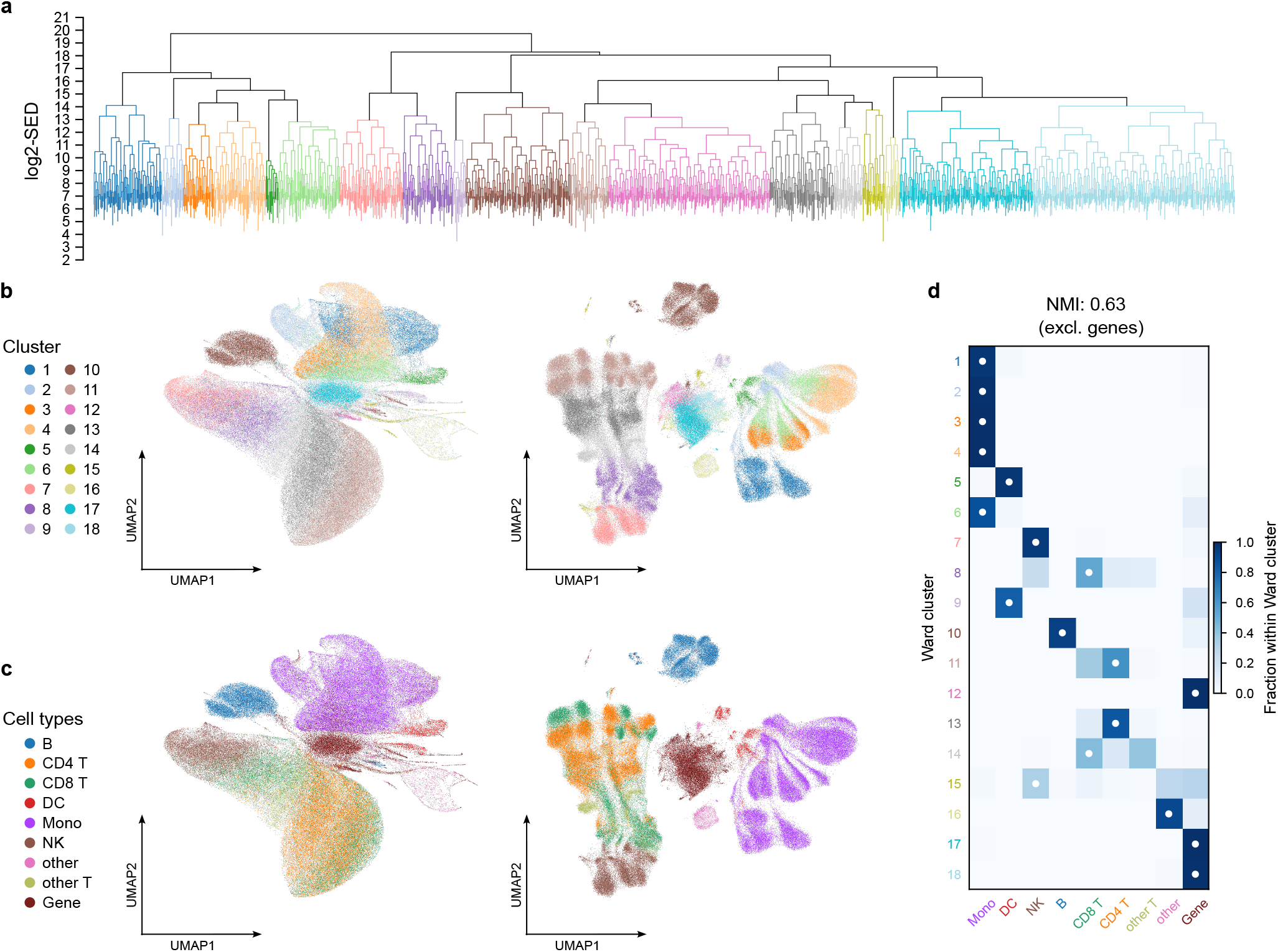
SCENE recovers a biologically structured joint cell–gene latent space in the PBMC CITE-seq dataset. **a**, Ward-linkage dendrogram of cells and genes in the learned latent space, with branches colored after cutting the dendrogram into log_2_(*N* + *G*) clusters for downstream comparison. **b**, UMAP visualization of the Laplacian initialization on the bipartite graph (left) and the converged SCENE latent representation (right), colored by Ward clusters computed in the converged latent space. **c**, The same representations as in **b**, colored by reference cell-type annotations, with genes shown as a separate node class. **d**, Correspondence between Ward clusters and reference annotations, shown as the fraction of each Ward cluster assigned to each cell-type or gene class.

### SCENE preserves cell identity while resolving perturbation responses

An informative representation of perturbed single-cell data should retain stable cell-type structure while capturing condition-specific transcriptional shifts. We assessed this trade-off in the IFN-*β* stim dataset [28], which spans diverse immune cell types and a strong IFN-*β* response. Fig. 3a,b shows that canonical marker genes occupy the expected cellular contexts in the joint latent space, with cells from the corresponding annotated populations generally lying at shorter Euclidean distances and higher predicted expression probabilities. This pattern is also observed for broadly expressed markers such as CD74, indicating that SCENE preserves cell-gene specificity while allowing genes to associate with multiple relevant cell populations. At the same time, SCENE resolves the IFN-*β* response as a distinct perturbation axis, canonical interferon-response genes localize to regions enriched for stimulated cells, separating IFN-*β*-treated cells from controls within the shared representation (Fig. 3c,d). These results show that SCENE retains cell identity and perturbation state as simultaneous biological axes within a shared cell-gene geometry.

**Figure 3.**
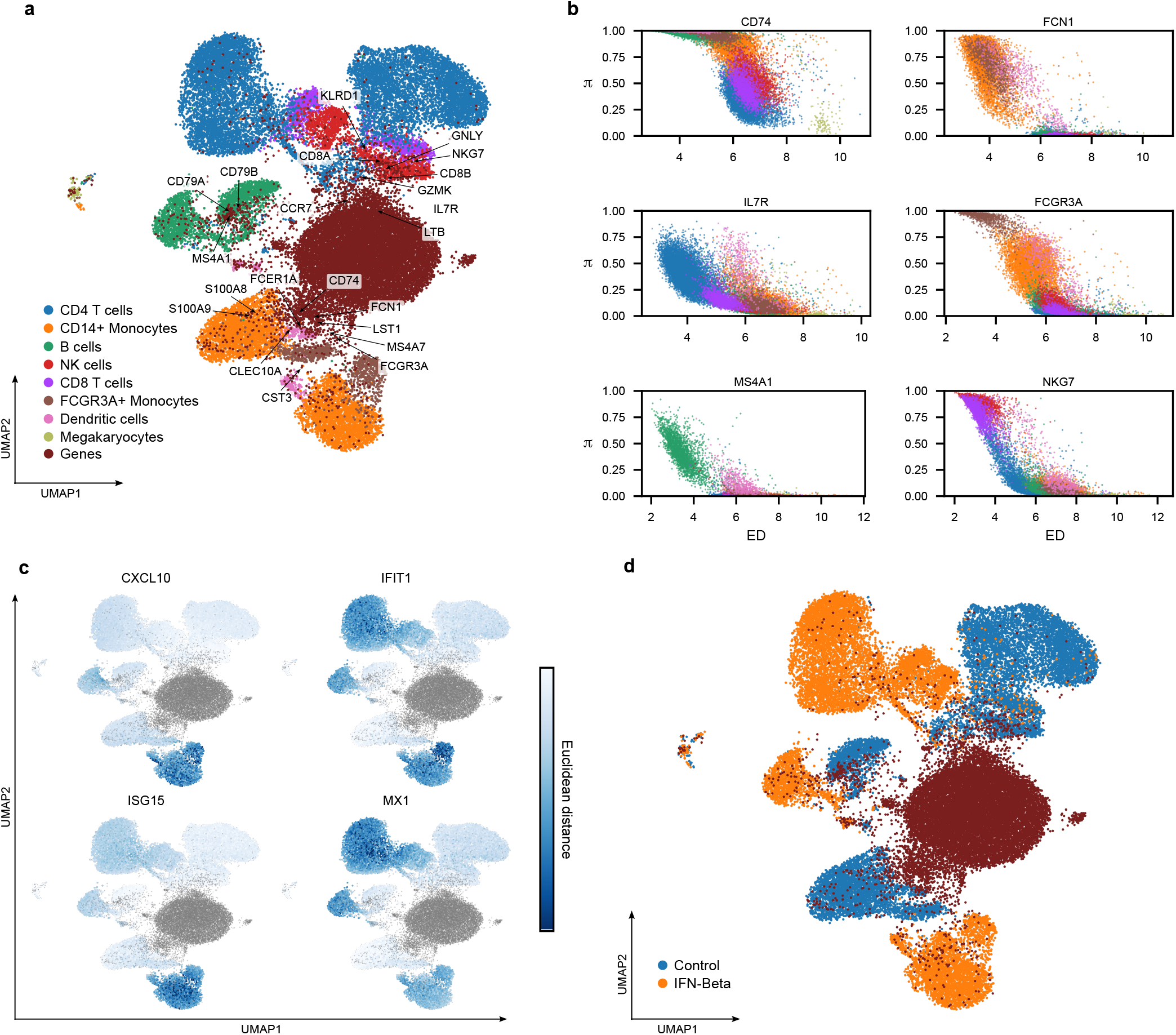
SCENE preserves cell identity while resolving IFN-*β* perturbation response in the joint cell–gene geometry of the IFN-*β* stim dataset. **a**, UMAP of the joint latent representation colored by cell type, with selected canonical marker genes indicating alignment between gene positions and known cellular identities. **b**, Predicted probability of gene expression as a function of Euclidean distance from selected canonical marker genes, with cells colored by annotation, showing that high-probability regions are enriched for the expected cell types. **c**, UMAP of the joint latent representation colored by Euclidean distance from cells to selected IFN-*β* response genes, revealing localization of perturbation-responsive programs within specific cellular compartments. **d**, UMAP of the joint latent representation colored by stimulation condition, showing that the perturbation-associated structure in **c** corresponds to stimulated and control groups.

### SCENE achieves strong batch integration while retaining biological signal

Comparative analysis often requires integrating datasets generated across laboratories and institutions, therefore an effective integration method must remove technical variation without erasing underlying biological structure. We assessed this trade-off across real and simulated benchmark datasets with diverse batch structures. SCENE achieves competitive or superior performance across real and simulated datasets on both biological-conservation and batch-correction metrics (Fig. 4b, full benchmark values across datasets and random seeds are reported in Supplementary Table 1). Its advantage is most pronounced on the SIM2 dataset, where nested batch effects present a particularly challenging setting for integration. As shown in Fig. 4d, SCENE recovers all four simulated cell groups under nested batch structure, whereas scVI largely collapses the biological groups into a mixed representation. Notably, SCENE remains biologically interpretable even in its native low-dimensional representation. In the HCA Nuclei dataset, the integrated representation mixes cells from the Harvard and Sanger batches while preserving clear cell-type organization (Fig. 4a). In the native twodimensional SCENE embedding, canonical cardiac marker genes remain localized near their expected cell populations, further supporting the biological interpretability of the integrated space (Fig. 4c). These results show that SCENE can align cells across batches while maintaining biological cell-type structure and retaining an interpretable cell–gene geometry even in native two-dimensional representations.

**Figure 4.**
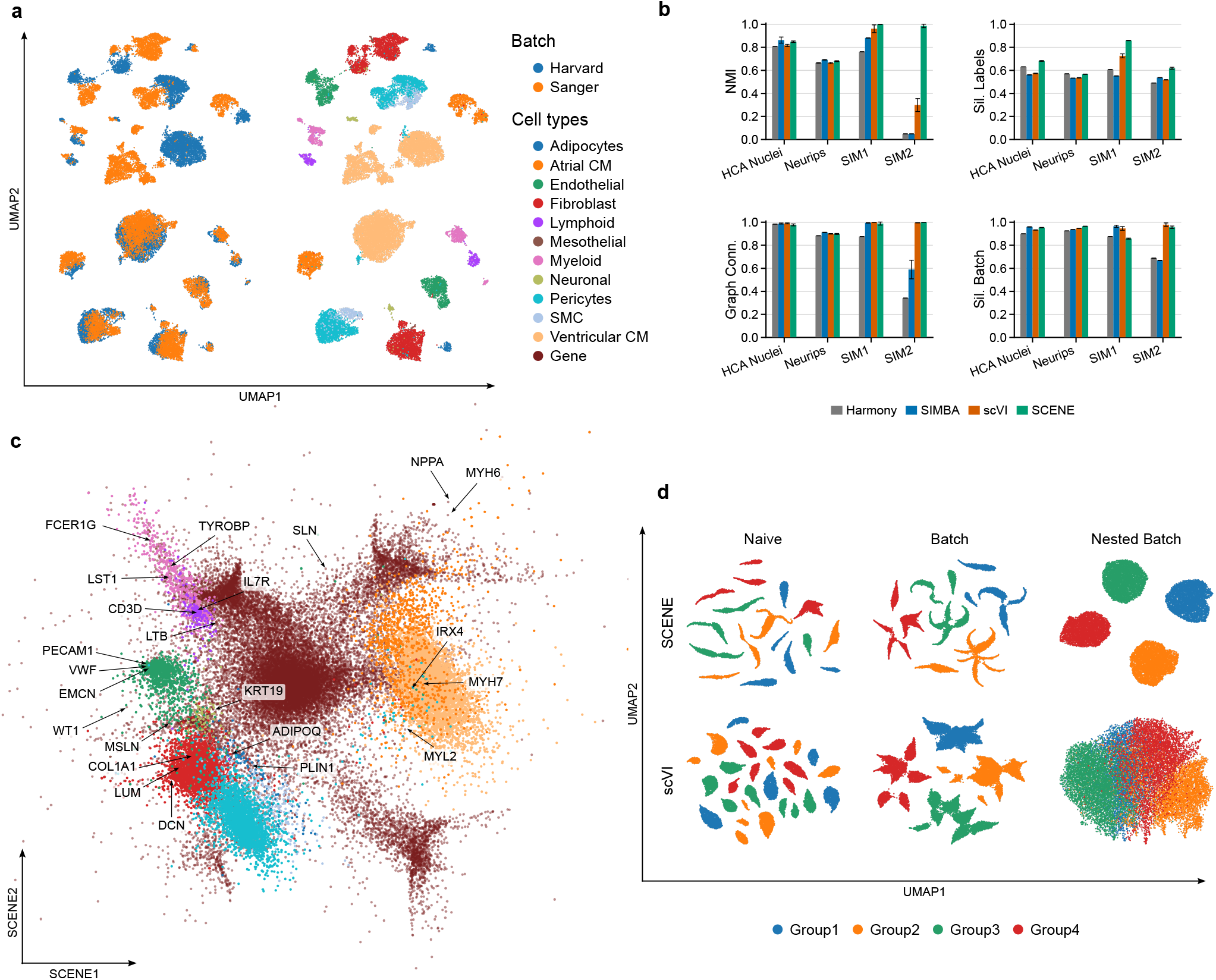
SCENE achieves strong batch integration performance while preserving biological structure. **a**, UMAP representations of cells from the HCA Nuclei dataset before integration (top) and after integration (bottom). **b**, Batch integration metrics comparing SCENE with Harmony, scVI, and SIMBA across multiple datasets. NMI and label silhouette measure biological conservation, whereas graph connectivity and batch silhouette measure batch correction performance. All metrics are oriented such that higher values indicate better performance. Error bars signify standard deviation across 5 runs with different seeds. **c**, Native 2D SCENE representation of the batch integrated HCA Nuclei dataset, showing canonical cardiac marker genes localized near their expected cell populations. **d**, Comparison of SCENE and scVI on the SIM2 dataset, illustrating improved performance of SCENE under nested batch effects.

### SCENE yields interpretable native ultra-low-dimensional embeddings

Many single-cell methods rely on post hoc visualization of higher-dimensional latent spaces, which can obscure whether biologically meaningful structure is truly present in the native representation. In contrast, SCENE yields interpretable ultra-low-dimensional embeddings while maintaining competitive predictive performance across latent dimensionalities (Fig. 5e).

**Figure 5.**
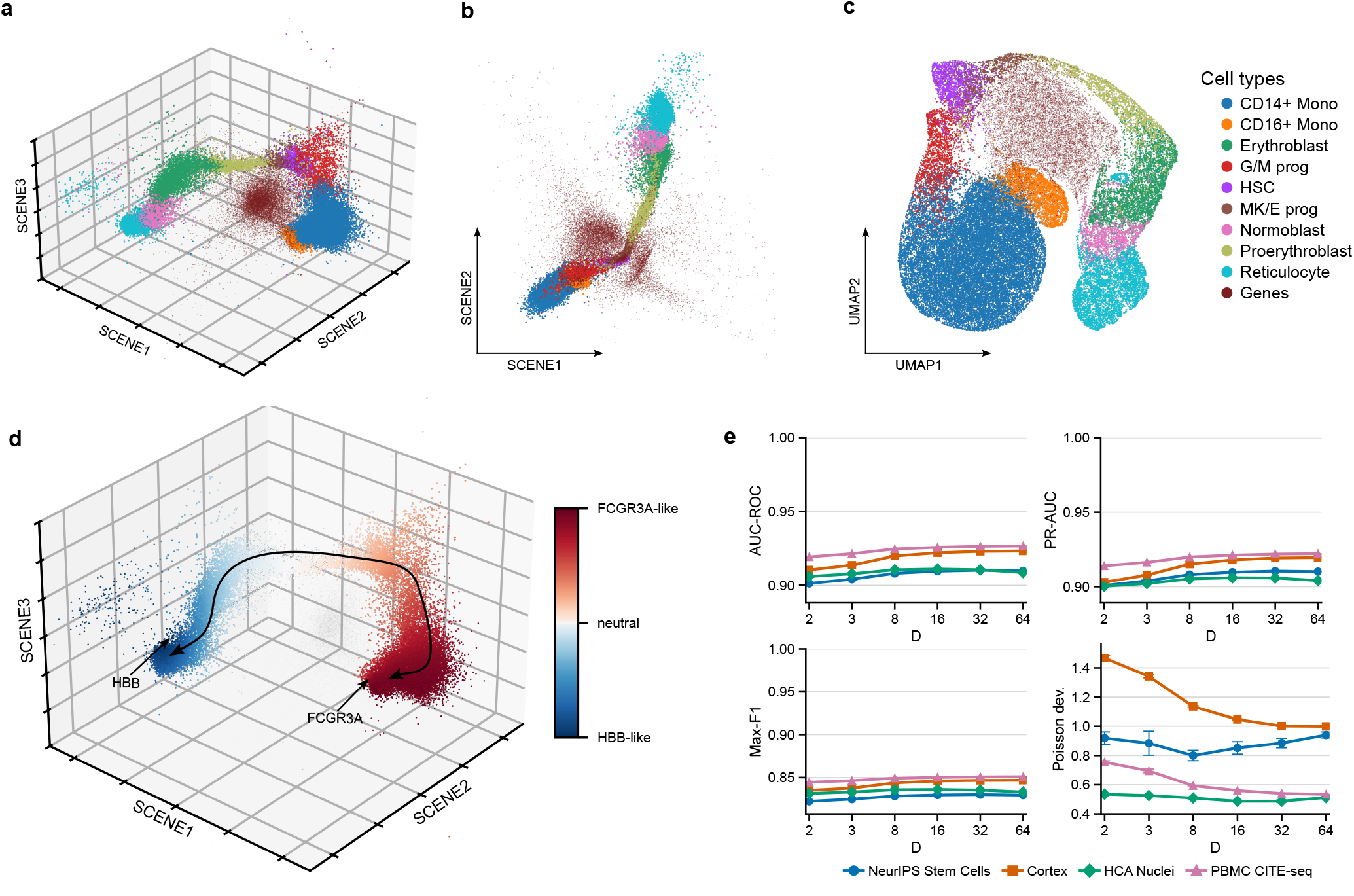
SCENE yields accurate and interpretable native ultra-low-dimensional embeddings. **a**, Native 3D latent representation of the stem cell subset from the NeurIPS dataset [29], showing that the two major lineage branches are clearly recovered in three dimensions. **b**, Native 2D latent representation of the same data, showing that the major lineage structure remains resolved even in two dimensions. **c**, A 2D UMAP of the native 3D representation shown in **a**, illustrating how post hoc visualization can alter the apparent global geometry of the learned representation. **d**, The same native 3D latent representation as in **a**, colored by relative Euclidean proximity to endpoint-associated genes for each lineage, showing that the latent geometry captures biologically meaningful cell–gene organization. **e**, Link prediction and count recovery performance across latent dimensionalities after masking a 10% held-out test set by setting its entries to zero. Link prediction performance saturates rapidly at low dimensionality, whereas count recovery remains more sensitive to dimensionality for some datasets.

Using a stem cell lineage subset from the NeurIPS dataset [29], we find that SCENE directly resolves monocytic and erythroid lineage branches from the common hematopoietic stem cell (HSC) in both three and two dimensions (Fig. 5a,b). The corresponding 2D UMAP of the native 3D representation alters the apparent global geometry (Fig. 5c), including changing the visual relationship between the erythroid and monocytic branch endpoints seen in the close proximity of reticulocytes and CD14+ monocytes. Moreover, endpoint-associated genes are positioned in biologically consistent regions of the joint cell-gene space, HBB aligns with the erythroid branch, whereas FCGR3A aligns with the monocyte branch (Fig. 5d). This indicates that the native geometry captures not only cell-cell lineage structure, but also interpretable cell-gene relationships.

To assess predictive performance across latent dimensionalities, we randomly selected 10% of nonzero entries and set their training counts to zero. These entries remained in the training matrix as zeros, while their original values were retained for evaluation. We evaluated recovery of their detection status and count magnitudes using link-prediction metrics and Poisson deviance, respectively (see Methods). Across datasets, link-prediction performance saturated rapidly at low dimensionality, whereas Poisson deviance continued to improve with additional dimensions for some datasets, demonstrating that very low-dimensional SCENE embeddings preserve much of the relational structure while higher dimensions mainly improve fine-grained count reconstruction (Fig. 5e). Notably for the stem cell subset in the NeurIPS dataset, dimensionality above 8D actually degraded count reconstruction performance. A side-by-side native 2D and 3D comparison with PCA, scVI and SIMBA is provided in Supplementary Fig. 1, and full link-prediction and count-recovery metrics across dimensions are reported in Supplementary Table 2.

### SCENE recovers biologically coherent perturbation programs

Just as biologically related cell types and states occupy coherent regions of the cell-cell latent geometry, genes participating in a common perturbation response may likewise organize into compact and biologically meaningful structures in gene space. To test this, we compared SCENE and SIMBA [18] in two perturbation settings with well-characterized biology, a glucocorticoid receptor (GR) stimulation time course present in the GR stim dataset and the stimulated CD4^+^ T cells from the TCR stim dataset.

In the first setting, we trained six independent embeddings for both SCENE and SIMBA on the GR breast line stimulation dataset [30] and examined how curated response modules were organized in gene space relative to a literature-based prior expectation (Fig. 6a). Module definitions and curation rationale are provided in Supplementary Table 3. Both models recovered aspects of the expected gene–gene organization, but SCENE produced a stronger and more consistent signal across different representations. This improvement was most evident for a module of canonical early direct GR target genes (GR Direct GRE, Supplementary Table 3) and a second module comprising glucocorticoid-induced transcription factors (TFs) expected to act later in the response (GR Late TF, Supplementary Table 3), which were recovered more robustly across the stimulation time course. The GR Direct GRE module showed greater relative compactness 18 hours after stimulation than in control cells (Fig. 6b,c), consistent with perturbation-associated organization of these genes in the learned latent space.

**Figure 6.**
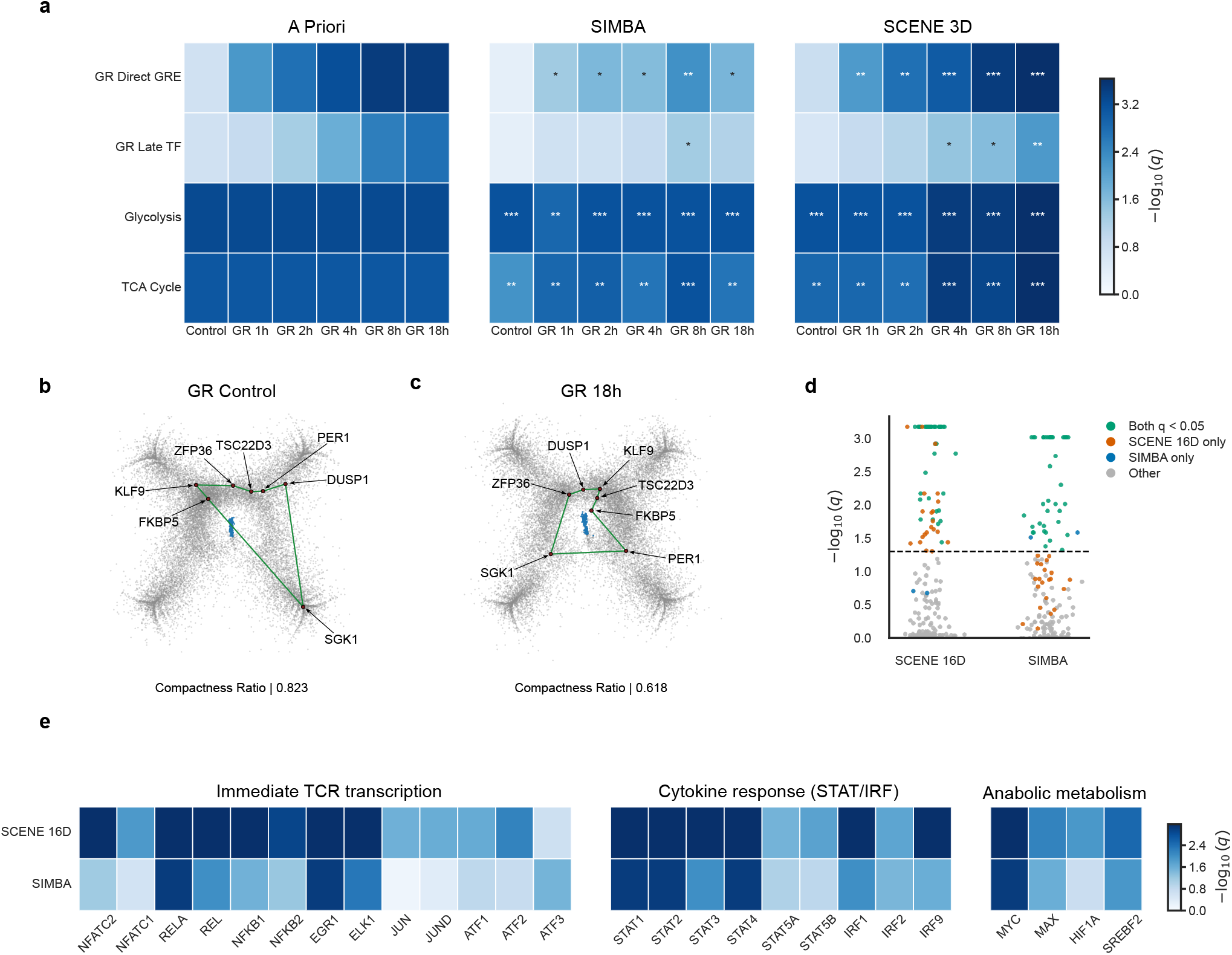
SCENE recovers expected perturbation programs in gene geometry better than SIMBA. **a**, Comparison of SIMBA and SCENE 3D in recovering expected gene responses to glucocorticoid receptor (GR) stimulation in the GR breast line stimulation dataset [30] across six conditions, with SCENE and SIMBA each trained separately for every condition. Statistical significance reflects the compactness of curated gene modules in latent space as a proxy for coordinated regulation, with each module evaluated against an empirical null distribution of 10,000 randomly sampled modules of matched size. **b**, Native 2D SCENE representation of GR control cells with genes from the GR Direct GRE module highlighted. **c**, Native 2D SCENE representation of GR-stimulated cells at 18 h with genes from the same GR Direct GRE module highlighted, showing increased compaction around the cells in the stimulated state. **d**, Comparison of SCENE and SIMBA in recovering transcription factor modules from DoRothEA confidence classes A and B. **e**, Heat map summarizing the recovery of 26 DoRothEA transcription factors in the CD4+ T-cell TCR stimulation dataset [31]. Control and stimulated cells were embedded jointly in the same latent space, and transcription factors were grouped into three TCR-relevant categories to illustrate recovery of response programs across multiple activation axes. Full transcription-factor module results are provided in the Supplementary Files. *q*: Benjamini–Hochberg-adjusted empirical *P* value; * *q <* 0.05, ** *q <* 0.01, *** *q <* 0.001. A Priori colors illustrate expectations, not significance.

In the second setting, we jointly embedded control and CD3/CD28-stimulated CD4+ T cells from the CD4+ T-cell TCR stimulation dataset [31] in a shared latent space to test whether the models could resolve a complex cellular activation response composed of many functionally distinct transcriptional events, while organizing these events coherently within the latent gene-space geometry. Using transcriptionfactor regulons from DoRothEA high confidence classes A and B [32], SCENE showed stronger and more consistent recovery of biologically coherent TF modules than SIMBA (Fig. 6d,e). In particular, SCENE more consistently resolved immediate-early TCR-associated transcriptional regulators, including NFATC1/2, RELA, REL, NFKB1/2, EGR1, ATF1/2 and ELK1, as well as cytokine-response regulators in the STAT/IRF axis and metabolic regulators such as MYC, MAX, HIF1A, and SREBF2. These results suggest that SCENE not only detects perturbation-responsive transcription factors, but also organizes them into biologically coherent functional programs within the latent geometry.

## Discussion

SCENE provides an interpretable probabilistic representation of scRNA-seq data by jointly embedding cells and genes in a shared Euclidean latent space directly from raw UMI counts. By combining a zero-inflated Bernoulli– Poisson likelihood with a bipartite latent-distance formulation, SCENE models sparse count structure while preserving an explicit geometric interpretation of cell–cell, cell–gene, and gene–gene relationships. This distinguishes SCENE from many graph-based single-cell methods, which typically construct graphs only after normalization, feature selection, or dimensionality reduction [15, 16, 19], and from encoder-based models, where useful latent representations are learned but the relationship between model parameters, genes, and cells is less directly interpretable [17, 33, 20, 21]. In SCENE, the observed count matrix itself defines the weighted bipartite graph, allowing representation learning to proceed directly from the original cell-gene observations rather than from a preprocessed surrogate.

Across multiple datasets and analysis settings, SCENE recovered biologically structured representations while remaining competitive with established single-cell methods. In benchmark integration tasks, SCENE matched or exceeded competing methods on biological-conservation and batch-correction metrics commonly used for single-cell integration evaluation [10], with particularly strong performance on simulated data with nested batch effects. Masked-count imputation on the PBMC CITE-seq dataset further supports this interpretation, with SCENE recovering held-out nonzero counts with lower reconstruction error and higher Pearson correlation than scVI (Supplementary Fig. 2).

SCENE embeds the geometry of the bipartite cell–gene graph into a latent space. Similar to recent efforts to co-embed cells and features [18, 19], we show that this geometry can be used not only for biologically coherent cell–cell relations but also for cell–gene and gene–gene relations. When inspecting relative cell–gene placement, we discover that known and specific marker genes position themselves close to their respective cell groups, validating that cell–gene distances in the latent representation of SCENE carry biological signal. In many single-cell workflows, latent representations are used primarily to organize cells for clustering, integration, or visualization, while biological interpretation is performed by returning to the original or normalized expression matrix [14, 4]. SCENE instead supports analysis directly in the learned latent representation. Because cells and genes occupy the same model-based Euclidean space, marker localization, cell-gene proximity, and gene-gene program structure can be evaluated within the representation used by the model itself.

The gene-space analyses extend this interpretation from cell identity to coordinated perturbation biology. In the GR stimulation time course, glucocorticoid-responsive genes became more compact according to their expected response dynamics, with early target-gene responses and later transcriptional-regulatory programs showing temporally distinct organization [30, 34]. Core metabolic reference programs, by contrast, remained coherently organized across the time course, indicating that SCENE can distinguish stable pathway-level coupling from dynamically induced perturbation responses. TCR stimulation analyses provided a complementary test of whether SCENE could organize latent gene geometry during a complex cellular response, where immediate-early transcriptional activation, cytokine-response circuitry, and metabolic remodeling unfold in parallel [35, 36]. Rather than collapsing these processes into a single activation signal, SCENE organized them as distinct but related functional neighborhoods within the same broader stimulated state. These results indicate that the latent gene geometry captures state-dependent functional coupling between genes, supporting direct distance-based biological analysis in the native SCENE space.

A central implication of these results is that SCENE’s native ultra-low-dimensional spaces can be inspected directly as model representations, rather than treated only as inputs to downstream visualization. Because the learned space is Euclidean and explicitly parameterized, distances between cells, genes, and cell-gene pairs retain their model-based interpretation in two or three dimensions. In the stem-cell lineage analysis (Fig. 5), SCENE clearly resolved the major erythroid and monocytic lineage branches directly in native 2D and 3D, while positioning endpoint-associated genes such as HBB and FCGR3A near their corresponding cellular trajectories. This makes the embedding directly interpretable as biological structure, marker-gene localization, and lineage relationships can be visually examined in the same coordinate system in which the model is fitted, without requiring an additional projection step. While there is a trade-off for performance we show that linkprediction saturates rapidly at low dimensionalities which further suggests that much of the relational structure of the cell–gene graph is captured by these compact native embeddings, whereas higher dimensions mainly improve fine-grained count reconstruction.

Although we focus here on scRNA-seq, the general formulation suggests a path toward other count-based singlecell modalities, including scATAC-seq peak counts and CITE-seq ADT counts. Because SCENE models an observed matrix as a weighted bipartite graph with a count likelihood, the framework is not inherently restricted to gene-expression measurements. This contrasts with many existing single-cell models, whose likelihoods, encoders, or graph constructions are designed around a particular molecular modality or preprocessing pipeline [17, 16, 19]. At the same time, extension to other modalities will likely require modality-specific likelihoods or random-effect structures. Extending SCENE to multimodal settings is therefore a natural direction, but should be evaluated empirically rather than assumed from the scRNA-seq results alone.

Scalability remains a practical concern for scRNA-seq analysis. SCENE defines an entry-wise likelihood over the cell–gene count matrix, so a dense implementation contains *O*(*NG*) likelihood terms, where *N* is the number of cells and *G* is the number of genes. However, this worst-case expression should be interpreted in the context of scRNA-seq data, *G* is bounded by the assayed gene set and does not grow with the number of cells. For a fixed feature space, increasing dataset size therefore primarily increases the cost linearly in *N*, with *G* acting as a dataset-dependent constant; in typical scRNA-seq workflows this feature space is on the order of 10^4^ genes, ranging from approximately 20,000 protein-coding genes to larger filtered transcriptomic reference sets [37]. In this paper, we evaluate the likelihood over the dense cell–gene graph at each training step, but the implementation also provides a cell-based mini-batching routine that restricts each update to a subset of cells and their associated gene-count profiles. More structural approximations, including hierarchical or block-distance extensions such as HBDM [26], may further reduce cost for very large graphs while preserving the latent-distance interpretation.

Evaluating performance presents inherent limitations in scRNA-seq, particularly within the standard benchmarking process. Benchmarks on real dataset annotations should be interpreted as proxies of true performance, since these mostly rely on reference annotations originally derived from other computational predictions rather than ground truth [38]. This further motivates the inclusion of simulated data, where the underlying structure is known, albeit simplified relative to real biological complexity [39].

SCENE provides a probabilistic framework for scRNA-seq data that jointly embeds cells and genes in a shared Euclidean latent space by modeling the count matrix as a weighted bipartite graph. Across diverse datasets, SCENE matches or exceeds established methods on cell-level representation and integration tasks while uniquely extending interpretability to cell-gene and gene-gene relations that reflect context-specific biological signals. By avoiding deep parameterizations and extensive preprocessing, SCENE provides a transparent metric space in which biological structure can be inspected directly. This structure is retained even in native two- and three-dimensional SCENE representations, making the interpretable model representations directly inspectable with meaningful cell–cell, cell– gene, and gene–gene geometry. These properties make SCENE a principled and extensible foundation for interpretable single-cell analysis, particularly for health and disease studies and for future applications to other singlecell count modalities.

## Methods

### Bipartite graph construction

SCENE directly models the count matrix as a network of cells and genes, expanding upon the high potential of graph-based single-cell methods [15, 16]. Specifically, we treat the count matrix 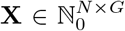 as the biadjacency matrix of a weighted bipartite graph *G* = (*V*_*c*_ ∪ *V*_*g*_, *ℰ*) where *V*_*c*_ and *V*_*g*_ are disjoint sets. Here *V*_*c*_ = {*c*_1_, …, *c*_*N*_} correspond to the set of cells with N nodes while *V*_*g*_ = {*g*_1_, …, *g*_*G*_} correspond to the set of genes with G nodes, hence an edge *e* ∈ ℰ can only occur between a cell and a gene. Since the network structure is analytically derived from **X**, each entry *x*_*ij*_ serves as the weight of the edge from cell *i* to gene *j*.

### Latent Distance Model

Our aim is to learn a joint, low-dimensional embedding of cells and genes without any preprocessing of the UMI counts. We build on the latent distance model (LDM) introduced by Hoff et al. in 2002 [25] which embeds each node *v* ∈ *V*_*c*_ ∪ *V*_*g*_ as a vector in a latent space ℝ^*d*^. As such, the bipartite approach [26] results in a joint low-dimensional latent embedding of both sets, i.e. a joint latent representation of cells and genes. Then let **z**_*i*_ ∈ ℝ^*d*^ denote the latent position of cell *i* for *i* = 1, …, *N* and **w**_*j*_ ∈ ℝ^*d*^ denote the latent position of gene *j*, for *j* = 1, …, *G*, in a latent space of dimensionality *d*.

The LDM assumes that the probability of an edge between a cell and a gene depends on the Euclidean distance in ℝ^*d*^, i.e., the closer two nodes are, the higher the probability of an edge. In our weighted bipartite graph, this distance further determines the edge weight, translating into the expression magnitude of a gene for a given cell.

As scRNA-seq UMI is inherently count data, we employ a probabilistic approach and model each cell-gene relation *x*_*ij*_ as arising from a Poisson distribution:

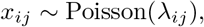

where the rate parameter is defined as in Nakis et al. [26] with node specific random effects:

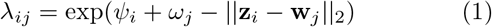

Here, *Ψ*_*i*_ and *ω*_*j*_ denote celland gene-specific random effects that absorb technical noise and batch effects separately from the latent variables **z**_*i*_ and **w**_*j*_, which are intended to encode genuine biological variability [7, 9].

Let 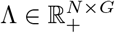 denote the matrix with entries *λ*_*ij*_. The log-likelihood under the above model can be expressed as

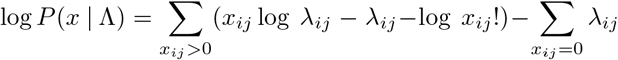

The Poisson assumption relies on E [*X*_*ij*_] = Var(*X*_*ij*_) = *λ*_*ij*_ and we presently model each cell-gene interaction *x*_*ij*_ as independent Poisson processes with interaction specific rates given by *λ*_*ij*_. This results in a distribution across all entries in **X** which can exhibit substantial overdispersion at population level. In other words, by modeling each observation with an individualized Poisson parameter, the model accommodates the excess variance typically observed in scRNA-seq data due to biological and technical variability [7, 9]. Other models instead use a negative-binomial likelihood and typically learn a shared dispersion parameter per gene (or per gene and cell group), alongside cell-gene specific means [33, 17, 27].

Nevertheless, the empirical distribution of counts in scRNA-seq is often marked by an abundance of zeros that far exceeds what is expected even under this heterogeneous Poisson model [7, 9]. Such sparsity reflects both biological absence and technical dropouts, and motivates the introduction of an explicit zero-inflation component [40, 11].

### Zero-inflated Poisson likelihood

To account for excess zeros, dropouts, and low-capture events, we use a Poisson hurdle formulation, in which a Bernoulli component models whether a cell-gene count is zero or positive, and a zero-truncated Poisson component models the positive count conditional on detection [41, 42].

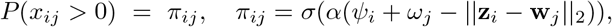

where *σ*(·) is the sigmoid function and *α >* 0 is a learnable scaling parameter which adapts the scale to the Bernoulli probability representation *π*_*ij*_ whereas the observed count values follow a truncated Poisson distribution in which the occurrence of zeros correspondingly has been removed. The full zero-inflated model then becomes:

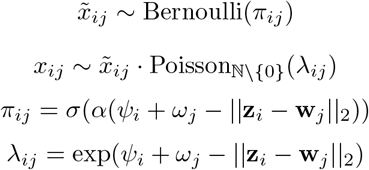

The truncated Poisson PMF is obtained by conditioning the standard Poisson distribution on the event that the observed count is strictly positive. For a Poisson random variable *X* ~ Poisson(*λ*_*ij*_), the standard PMF is given by 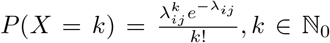. Since zero counts are modeled separately through the Bernoulli component, the Poisson distribution must be re-normalized over the support ℕ \ {0}. This yields

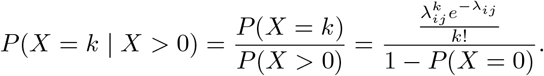

Using that

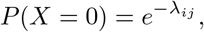

we obtain

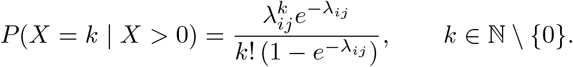

Resulting in the PMF:

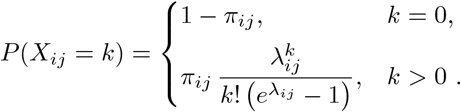

Under this likelihood, the learned probability that gene *j* is expressed in cell *i* is given directly by *π*_*ij*_. The corresponding expected observed count is obtained from the PMF as

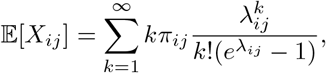

and using

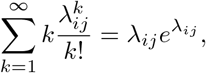

this becomes

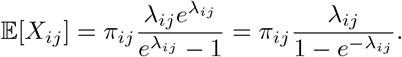

Thus, *λ*_*ij*_ is the underlying Poisson rate parameter, whereas E[*X*_*ij*_] is the model-implied expected count. Notably, conditioned on expression, the expected positive count becomes:

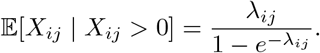

Finally, let Π ∈ [0, 1]^*N×G*^ denote the matrix of nonzero-expression probabilities, with entries *π*_*ij*_. The total zero-inflated log-likelihood is then given by

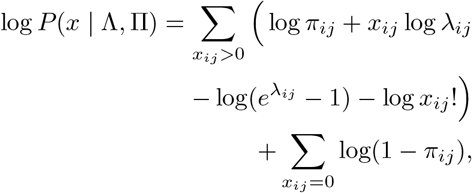

All model parameters (*α, Ψ*_*i*_, *ω*_*j*_, **z**_*i*_, **w**_*j*_) are learned by maximizing the total log-likelihood with the AdamW optimizer [43]. Weight decay of 10^*−*3^ is applied to the latent cell and gene embeddings, while no weight decay is applied to the cell and gene random effects or the trainable parameter underlying *α*.

We evaluate the empirical behavior of this zero-inflated formulation using held-out count recovery and detection-intensity calibration diagnostics in Supplementary Figs. 3 and 4.

### Explicit modeling of known batch effects

We extend SCENE to account for explicit batch effects, a central challenge in scRNA-seq [10] that is also addressed by methods such as Harmony [23], Scanorama [24], and scVI [17]. Rather than correcting the count matrix in a preprocessing step, SCENE includes known batch effects directly in the likelihood through additive batch-specific terms in the cell-gene interaction.

Let *b*_*l*_(*i*) ∈ {1, …, *B*_*l*_} denote the batch assignment of cell *i* for batch layer *l*, where *B*_*l*_ is the number of batches in that layer and *l* = 1, …, *L*. We extend the predictors for *λ*_*ij*_ and *π*_*ij*_ as

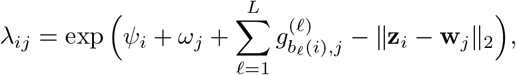

and

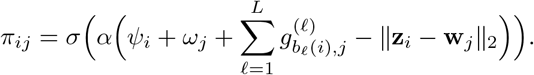

Here, *Ψ*_*i*_ and *ω*_*j*_ are cell- and gene-specific random effects, respectively, while 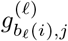 denotes the gene-specific batch adjustment for gene *j* in the batch assigned to cell *i* considering batch layer *l*. Thus, cells assigned to the same batch receive the same gene-specific correction profile, while different genes may be affected differently by the same batch.

For each batch layer *l*, we consider two possible parameterizations of 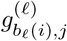. In the full parameterization, each batch-gene combination has its own free parameter,

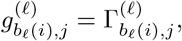

where

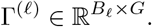

This gives *B*_*l*_*G* additional parameters for batch layer *l*, where *G* is the number of genes. This formulation is the most flexible because it allows each batch to have an unrestricted gene-specific effect. In the low-rank parameterization, we define

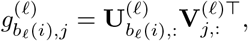

Where

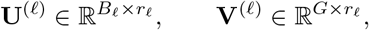

and *r*_*l*_ is the chosen rank for batch layer *l*. This gives

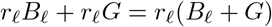

additional parameters. The low-rank formulation imposes shared structure across genes and batches, reducing the number of parameters while still allowing batch effects to vary across genes. The batch terms are estimated jointly with the latent cell and gene embeddings, allowing SCENE to separate known technical variation from the biological signal encoded by **z**_*i*_ and **w**_*j*_. All parameters for batch effects are also regularized in the AdamW optimizer using a weight decay of 10^*−*3^.

### Initialization using the Graph Laplacian

We introduce an initialization approach based on a spectral embedding of the weighted bipartite graph induced by the count matrix. This approach can be used to initialize the cell and gene latent positions. Let 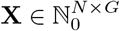 denote the cell-gene biadjacency. Define degree vectors and the block adjacency and degree

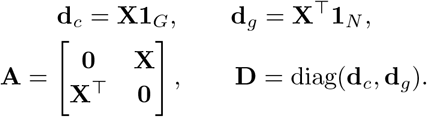

We use the symmetric normalized Laplacian as introduced in spectral graph theory [44].

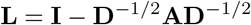

The initialization is then obtained from the non-trivial eigenvectors of **L**, following the Laplacian Eigenmaps approach [45]. Specifically, let **q**_1_, …, **q**_*k*_ be the *k* eigen-vectors associated with the smallest positive eigenvalues, excluding the trivial eigenvector associated with eigenvalue 0. Stacking these yields **Q** ∈ ℝ ^(*N* +*G*)*×k*^. We scale each row of **Q** by the inverse square root of the corresponding node degree to obtain the initial coordinates **Y** = **D**^*−*1*/*2^**Q**. We then partition **Y** by node type: the first *N* rows yield the initial cell embeddings **Z**_cells_, and the last *G* rows yield the initial gene embeddings **W**_genes_.

### Imputation

For any missing cell-gene pair we simply predict the observed count as 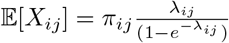. The bipartite graph couples every cell to every gene it expresses, so each gene constrains the placement of cells and vice-versa. This global property of mutual embedding allows dropout events and stochastic count fluctuations to be inferred directly from the latent geometry.

### Significance testing of gene-module compactness in latent gene space

To test whether predefined biological gene modules occupy unusually compact regions of the latent gene embedding, we evaluated each module against an empirical null distribution of random gene sets of matched size. Let **w**_*j*_ ∈ ℝ^*d*^ denote the latent embedding of gene *j*, and let *M* ⊂ {1, …, *G}* denote the set of genes belonging to a predefined module after intersecting the module with the genes present in the dataset.

For each module, we quantified its observed spatial compactness relative to the background gene space. We first computed the mean pairwise distance among genes in the module,

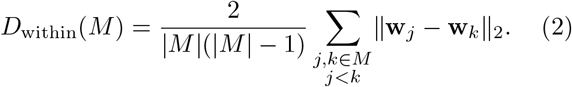

We then computed the mean distance between module genes and all genes outside the module,

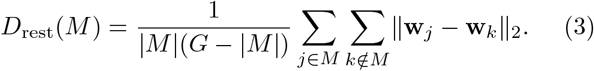

The test statistic was defined as the relative compactness score

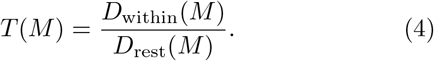

Lower values of *T* (*M*) indicate that genes in the module are closer to one another than expected relative to their average distance to the remaining genes.

Statistical significance was assessed by permutation. For each module *M*, we generated *B* = 10,000 random gene sets *M*_*b*_ by sampling genes uniformly without replacement from the full gene set, with |*M*_*b*_| = |*M* |. For each random gene set, we recomputed the same statistic *T* (*M*_*b*_). The empirical one-sided *P* value was calculated using the standard finite-sample correction [46]

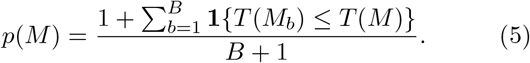

This test asks whether the observed module is more compact in latent gene space than expected for random gene sets of the same size. Empirical *P* values were adjusted across tested modules using the Benjamini–Hochberg false discovery rate procedure.

### Evaluation metrics

We use evaluation metrics targeting two aspects of model performance, held-out prediction of cell-gene observations and quality of the learned cell-level representation. Link prediction and Poisson deviance assess recovery of masked entries in the count matrix, while NMI, silhouette score, and graph connectivity assess clustering quality and integration performance of the cell embeddings.

### Link Prediction

We evaluated link prediction as recovery of held-out nonzero cell-gene interactions in the bipartite count graph. We held out 10% of nonzero entries by setting the corresponding training counts to zero. After optimization, each held-out cell-gene pair (*i, j*) was scored using the model-implied probability of nonzero expression. For SCENE-ZIP this score is given directly by

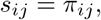

Held-out positive edges were evaluated against an equal-sized set of sampled zero entries from the same cell-gene matrix.

Performance was summarized using ROC-AUC, trapezoidal PR-AUC, and maximum positive-class F1 across thresholds. This evaluation tests whether the learned latent geometry generalizes beyond the observed training graph by assigning higher detection probability to missing true cell-gene links than to unobserved pairs.

### Poisson Deviance

To assess recovery of held-out count magnitudes, we computed the mean Poisson deviance on the held-out nonzero entries. For SCENE-ZIP, predicted counts were given by the zero-truncated Poisson mean,

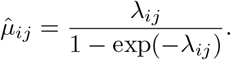

For the held-out set *S*_test_, mean Poisson deviance was computed as

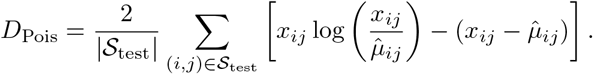

Lower values indicate better recovery of held-out count magnitudes.

### Normalized Mutual Information (NMI)

NMI measures the similarity between predicted clusters *U* and reference labels *V*. It is defined as:

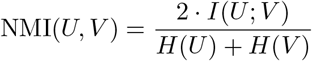

where *I*(*U*; *V*) is the mutual information and *H*(·) denotes entropy. Values lie in [0,1], with 1 indicating perfect agreement between clusters and labels [47].

### Silhouette Score

For each sample *i*, let *a*(*i*) denote the mean intra-cluster distance, average distance between i and all other points in its cluster, and let *b*(*i*) denote the mean nearest-cluster distance (minimum average distance between *i* and points in any other cluster). The silhouette score for sample *i* is

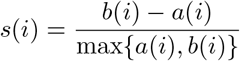

The global score is the average *s*(*i*) across all samples. Scores close to 1 indicate well-separated clusters, scores near 0 indicate overlapping clusters, and negative scores suggest misclassification [48]. For label silhouette, we rescaled the mean score to [0, 1] as 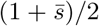. For batch silhouette, we computed the mean silhouette using batch labels within each cell type containing at least three cells and two batches, transformed it as 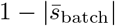, and averaged across eligible cell types weighted by their cell counts. All silhouettes used Euclidean distances. Higher transformed scores indicate stronger cell-type separation or weaker within-cell-type batch separation, respectively.

### Graph Connectivity

For each cell type *c*, let *C* denote the set of cells assigned to *c*, and let *LCC*(*C*) denote the largest connected component of *C* in the kNN graph. The connectivity score is

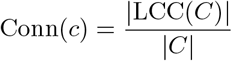

The overall graph connectivity is the mean of Conn(*c*) across all cell types. A score of 1 indicates that all cells of a given type are connected in the embedding space, while lower values indicate fragmentation across batches [10].

### Datasets

We evaluated SCENE on nine different scRNA-seq and Single-nucleus RNA sequencing (snRNA-seq) datasets, two of which are simulated, while the remaining seven are from actual samples spanning multiple tissues, experimental conditions and sequencing technologies.

#### Cortex

scRNA-seq dataset of 3,005 cells from the mouse somatosensory cortex and hippocampus, generated using the Fluidigm C1 platform with UMI-based transcript counting [49].

#### HCA Nuclei

scRNA-seq and snRNA-seq dataset of ~ 487,000 cells and nuclei from six regions of the adult human heart, capturing major cardiac and stromal cell types with droplet-based technologies [50], we use a subset of 13,733 snRNA cells.

#### PBMC CITE-seq

scRNA-seq dataset of 152,094 peripheral blood mononuclear cells (PBMC) from eight human donors, collected before and after HIV vaccination, originally generated as part of a multimodal CITE-seq atlas with paired protein measurements [12].

#### SIM1

Simulated scRNA-seq dataset of 12,097 cells across 6 batches, generated using the Splatter package to represent variation in cell-type compositions across batches for benchmarking integration methods [10].

#### SIM2

Simulated scRNA-seq dataset of 19,318 cells across 16 batches and 4 cell groups, generated with the Splatter package to model nested batch effects with cell-type compositional variation for benchmarking integration methods [10].

#### IFN-*β* stim

scRNA-seq dataset of PBMCs from eight lupus patients, split into untreated and recombinant IFN-*β*stimulated conditions for 6 h, generated using droplet-based scRNA-seq with sample multiplexing by natural genetic variation [28].

#### NeurIPS

multimodal single-cell dataset of human bone marrow mononuclear cells generated for the NeurIPS 2021 benchmark on prediction and integration of DNA, RNA, and protein measurements in single cells. We use the RNA expression component, including a hematopoietic stem and progenitor cell lineage subset spanning erythroid and monocytic differentiation trajectories [29].

#### GR stim

scRNA-seq dataset of T47D A1–2 human breast cancer cells treated with dexamethasone for 1, 2, 4, 8, and 18 h, with vehicle-treated cells as controls, generated using the Bio-Rad ddSEQ/SureCell platform [30].

#### TCR stim

scRNA-seq dataset of purified human CD4+ T cells from PBMCs, either unstimulated or stimulated for 4 h with anti-CD3/CD28 Dynabeads, generated using the 10x Genomics Chromium Single Cell 3’ v2 platform [31].

### Data quality control

For all analyses, we used publicly available curated count matrices as distributed by the original studies or benchmark resources. Before model fitting, we performed a quality-control audit to verify that the matrices were suitable for direct count modeling. For each dataset, we inspected library sizes, the number of detected genes per cell, the presence of zero-count cells or cells with no detected genes, feature-level detection rates, and mitochondrial fractions when gene identifiers allowed mitochondrial genes to be identified.

## Supporting information

Supplementary Data - SCENE TCR

Supplementary Data - SIMBA TCR

## Data availability

All datasets used in this study are publicly available from previously published sources. The mouse cortex/hippocampus UMI count matrix was loaded with the scvi-tools cortex() loader, which provides the Zeisel et al. dataset associated with the Linnarsson laboratory cortex resource and GEO accession GSE60361 [49]. The Heart Cell Atlas nuclei dataset was derived from the Litvinukova et al. Heart Cell Atlas resource through the scvi-tools heart cell atlas subsampled() loader [50]. The PBMC CITE-seq RNA count data were loaded with the scvi-tools pbmc seurat v4 cite seq() loader and are derived from the Seurat v4 multimodal PBMC reference from Hao et al., with RNA counts associated with GEO accession GSE164378 [12]. The SIM1 and SIM2 simulated datasets were obtained from the benchmark matrices used by Luecken et al. [10]. The IFN-*β* stimulation dataset was loaded through the pertpy kang 2018() curated copy of the Kang et al. PBMC stimulation dataset [28]. The NeurIPS bone marrow CITE-seq dataset was obtained from the OpenProblems/NeurIPS 2021 processed file associated with GEO accession GSE194122 [29]. The glucocorticoid receptor stimulation dataset was obtained from Hoffman et al. through GEO accession GSE141834 [30]. The TCR stimulation dataset was obtained from Ding et al. through GEO accession GSE147928 [31].

## Code availability

The SCENE package is available on PyPI at https://pypi.org/project/scene-ldm, with source code and installation instructions at https://github.com/oscarmoeberg/SCENE. Analysis scripts and environment specifications for reproducing the results reported in this manuscript are available at https://github.com/oscarmoeberg/SCENE-reproducibility.

## Supplementary Tables

### Benchmark and reconstruction metrics

**Supplementary Table 1.** Benchmark metrics comparing SCENE to Harmony, SIMBA, and scVI across datasets. Entries are mean *±* standard deviation across five seeds. Bold indicates the best mean within each dataset and metric; underlining indicates the second-best mean. Dagger denotes datasets with batch correction.

| Dataset | Method | Bio-conservation |  | Batch-correction |  |
| --- | --- | --- | --- | --- | --- |
|  |  | Sil. labels | NMI | Sil. batch | Graph conn. |
| Cortex | PCA | <u>0.611</u> ± 0.000 | 0.692 ± 0.000 | - | - |
| | SIMBA | 0.549 ± 0.001 | <u>0.778</u> ± 0.030 | - | - |
| | scVI | 0.600 ± 0.001 | 0.748 ± 0.005 | - | - |
| | SCENE | **0.627** ± 0.001 | **0.848** ± 0.004 | - | - |
| PBMC CITE-seq | PCA | **0.571** ± 0.000 | <u>0.657</u> ± 0.000 | - | - |
| | SIMBA | 0.544 ± 0.000 | **0.675** ± 0.004 | - | - |
| | scVI | 0.535 ± 0.003 | 0.610 ± 0.006 | - | - |
| | SCENE | <u>0.562</u> ± 0.001 | 0.637 ± 0.004 | - | - |
| HCA Nuclei<sup>†</sup> | Harmony | <u>0.631</u> ± 0.000 | 0.807 ± 0.000 | 0.900 ± 0.000 | 0.984 ± 0.000 |
| | SIMBA | 0.561 ± 0.001 | **0.863** ± 0.027 | **0.959** ± 0.002 | <u>0.988</u> ± 0.005 |
| | scVI | 0.574 ± 0.002 | 0.817 ± 0.009 | 0.933 ± 0.001 | **0.990** ± 0.004 |
| | SCENE | **0.682** ± 0.003 | <u>0.849</u> ± 0.007 | <u>0.954</u> ± 0.002 | 0.977 ± 0.008 |
| NeurIPS<sup>†</sup> | Harmony | **0.570** ± 0.000 | 0.665 ± 0.002 | 0.925 ± 0.000 | 0.883 ± 0.000 |
| | SIMBA | 0.533 ± 0.000 | **0.692** ± 0.003 | 0.936 ± 0.000 | **0.912** ± 0.002 |
| | scVI | 0.535 ± 0.002 | 0.664 ± 0.006 | <u>0.948</u> ± 0.001 | <u>0.900</u> ± 0.003 |
| | SCENE | <u>0.567</u> ± 0.001 | <u>0.680</u> ± 0.004 | **0.965** ± 0.001 | 0.899 ± 0.004 |
| SIM1<sup>†</sup> | Harmony | 0.607 ± 0.000 | 0.762 ± 0.000 | 0.877 ± 0.000 | 0.875 ± 0.000 |
| | SIMBA | 0.552 ± 0.001 | 0.882 ± 0.001 | **0.965** ± 0.009 | <u>0.995</u> ± 0.002 |
| | scVI | <u>0.726</u> ± 0.017 | <u>0.962</u> ± 0.034 | <u>0.946</u> ± 0.016 | **1.000** ± 0.000 |
| | SCENE | **0.859** ± 0.002 | **1.000** ± 0.000 | 0.858 ± 0.005 | 0.988 ± 0.013 |
| SIM2<sup>†</sup> | Harmony | 0.491 ± 0.000 | 0.049 ± 0.000 | 0.688 ± 0.000 | 0.342 ± 0.000 |
| | SIMBA | <u>0.537</u> ± 0.001 | 0.049 ± 0.000 | 0.668 ± 0.002 | 0.590 ± 0.081 |
| | scVI | 0.518 ± 0.002 | <u>0.299</u> ± 0.056 | **0.979** ± 0.015 | <u>0.996</u> ± 0.001 |
| | SCENE | **0.619** ± 0.008 | **0.986** ± 0.015 | <u>0.956</u> ± 0.011 | **1.000** ± 0.000 |

**Supplementary Table 2.** SCENE reconstruction and prediction metrics across latent dimensions for each dataset. Entries are mean *±* standard deviation across seeds. Bold indicates the best mean within each dataset and metric; underlining indicates the second-best mean. For Poisson deviance, lower is better.

| Dataset | Dim. | AUC-ROC | PR-AUC | Max F1 | Poisson Dev. |
| --- | --- | --- | --- | --- | --- |
| NeurIPS Stem Cells | 2 | 0.9013 ± 0.0003 | 0.9009 ± 0.0003 | 0.8221 ± 0.0002 | 0.9191 ± 0.0420 |
| | 3 | 0.9041 ± 0.0005 | 0.9039 ± 0.0005 | 0.8247 ± 0.0005 | 0.8838 ± 0.0821 |
| | 8 | 0.9081 ± 0.0002 | 0.9081 ± 0.0001 | 0.8283 ± 0.0002 | **0.7996** ± 0.0351 |
| | 16 | 0.9096 ± 0.0001 | 0.9097 ± 0.0001 | <u>0.8296</u> ± 0.0001 | <u>0.8518</u> ± 0.0427 |
| | 32 | **0.9102** ± 0.0000 | **0.9105** ± 0.0000 | **0.8300** ± 0.0000 | 0.8853 ± 0.0326 |
| | 64 | <u>0.9097</u> ± 0.0001 | <u>0.9101</u> ± 0.0001 | 0.8294 ± 0.0001 | 0.9401 ± 0.0206 |
| Cortex | 2 | 0.9104 ± 0.0003 | 0.9030 ± 0.0005 | 0.8350 ± 0.0003 | 1.4671 ± 0.0162 |
| | 3 | 0.9136 ± 0.0002 | 0.9077 ± 0.0003 | 0.8376 ± 0.0001 | 1.3419 ± 0.0111 |
| | 8 | 0.9198 ± 0.0002 | 0.9152 ± 0.0002 | 0.8437 ± 0.0003 | 1.1360 ± 0.0053 |
| | 16 | 0.9221 ± 0.0000 | 0.9180 ± 0.0000 | 0.8459 ± 0.0001 | 1.0467 ± 0.0034 |
| | 32 | <u>0.9230</u> ± 0.0000 | <u>0.9193</u> ± 0.0000 | <u>0.8466</u> ± 0.0000 | <u>1.0014</u> ± 0.0016 |
| | 64 | **0.9233** ± 0.0000 | **0.9196** ± 0.0000 | **0.8468** ± 0.0001 | **0.9992** ± 0.0005 |
| HCA Nuclei | 2 | 0.9060 ± 0.0002 | 0.9003 ± 0.0001 | 0.8313 ± 0.0002 | 0.5349 ± 0.0017 |
| | 3 | 0.9078 ± 0.0002 | 0.9022 ± 0.0002 | 0.8330 ± 0.0002 | 0.5256 ± 0.0019 |
| | 8 | 0.9105 ± 0.0001 | 0.9053 ± 0.0001 | <u>0.8356</u> ± 0.0001 | 0.5087 ± 0.0071 |
| | 16 | **0.9110** ± 0.0001 | **0.9060** ± 0.0002 | **0.8360** ± 0.0001 | **0.4868** ± 0.0026 |
| | 32 | <u>0.9105</u> ± 0.0000 | <u>0.9059</u> ± 0.0001 | 0.8352 ± 0.0001 | <u>0.4872</u> ± 0.0073 |
| | 64 | 0.9086 ± 0.0000 | 0.9041 ± 0.0000 | 0.8330 ± 0.0000 | 0.5117 ± 0.0022 |
| PBMC CITE-seq | 2 | 0.9193 ± 0.0004 | 0.9140 ± 0.0003 | 0.8445 ± 0.0002 | 0.7543 ± 0.0096 |
| | 3 | 0.9214 ± 0.0001 | 0.9164 ± 0.0002 | 0.8463 ± 0.0001 | 0.6942 ± 0.0129 |
| | 8 | 0.9247 ± 0.0001 | 0.9200 ± 0.0001 | 0.8492 ± 0.0001 | 0.5933 ± 0.0020 |
| | 16 | 0.9258 ± 0.0000 | 0.9212 ± 0.0000 | 0.8501 ± 0.0000 | 0.5599 ± 0.0007 |
| | 32 | <u>0.9264</u> ± 0.0000 | <u>0.9219</u> ± 0.0000 | <u>0.8507</u> ± 0.0000 | <u>0.5398</u> ± 0.0004 |
| | 64 | **0.9266** ± 0.0000 | **0.9222** ± 0.0000 | **0.8509** ± 0.0000 | **0.5338** ± 0.0002 |

## Supplementary Figures

**Supplementary Figure 1.**
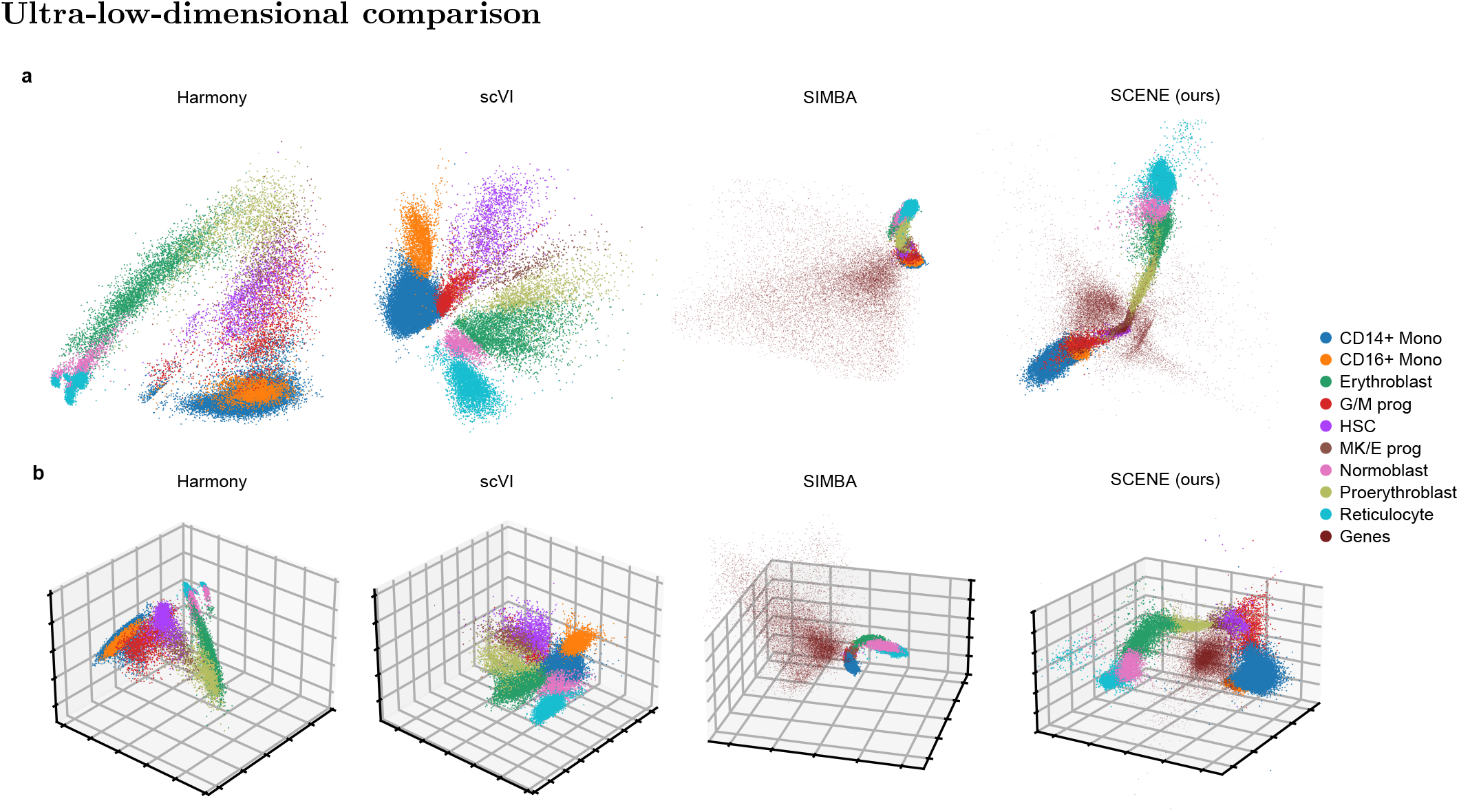
Ultra-low-dimensional comparison on the stem cell subset of the NeurIPS dataset. **a**, Native 2D representations of Harmony, scVI, SIMBA and SCENE. **b**, Native 3D representations of Harmony, scVI, SIMBA and SCENE. Across all models we correct for batches.

**Supplementary Figure 2.**
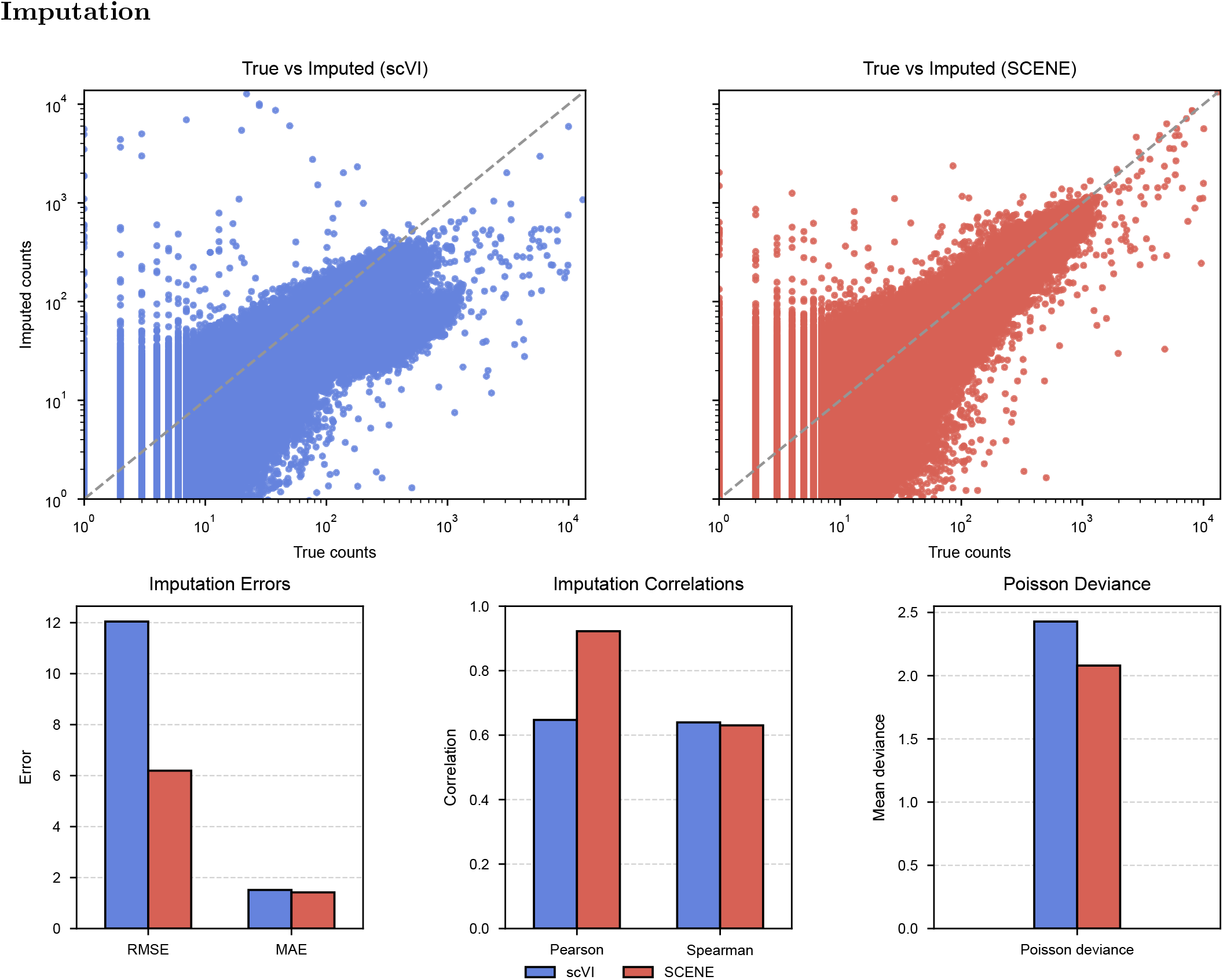
Imputation metrics for SCENE and scVI. Accuracy of gene-expression imputation on the PBMC CITE-seq dataset. A random 10% of nonzero counts were flipped to zero during training and subsequently predicted by each model. The top row shows observed counts versus imputed counts for scVI (left) and SCENE (right); the dashed line marks the identity. Bottom row shows aggregate performance on the held-out entries. The left panel reports root-mean-square error (RMSE) and mean absolute error (MAE); the middle panel shows Pearson and Spearman correlations between true and imputed counts: the right panel reports the Poisson deviance. SCENE achieves roughly half the reconstruction error of scVI and increases the linear correlation (Pearson *≈* 0.9) while also achieving a lower Poisson deviance, while both methods attain comparable rank concordance.

**Supplementary Figure 3.**
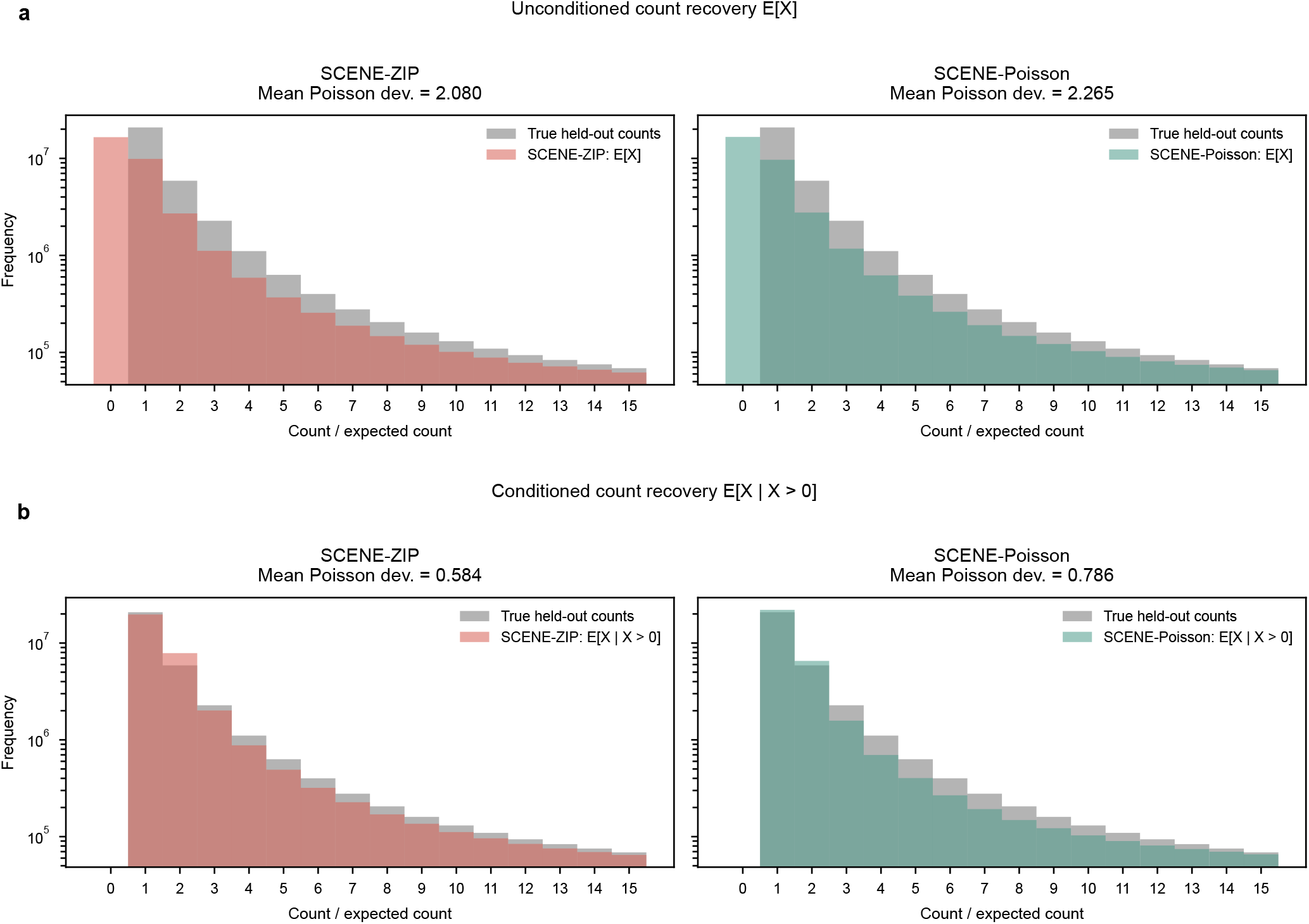
Zero-inflated likelihood improves held-out count recovery on the PBMC CITE-seq dataset. **a**, Comparison of empirical held-out count frequencies with model-implied expected counts under SCENE-ZIP and SCENE-Poisson. SCENE-ZIP better matches the full held-out count distribution and reduces mean Poisson deviance from 2.265 to 2.080, an 8.2% reduction. **b**, The same comparison conditioned on nonzero expression, using E [*X* | *X >* 0] for the zero-inflated model. Conditioning removes the explicit zero process and compares recovery of positive count magnitudes, where SCENE-ZIP reduces mean Poisson deviance from 0.786 to 0.584, a 25.7% reduction.

**Supplementary Figure 4.**
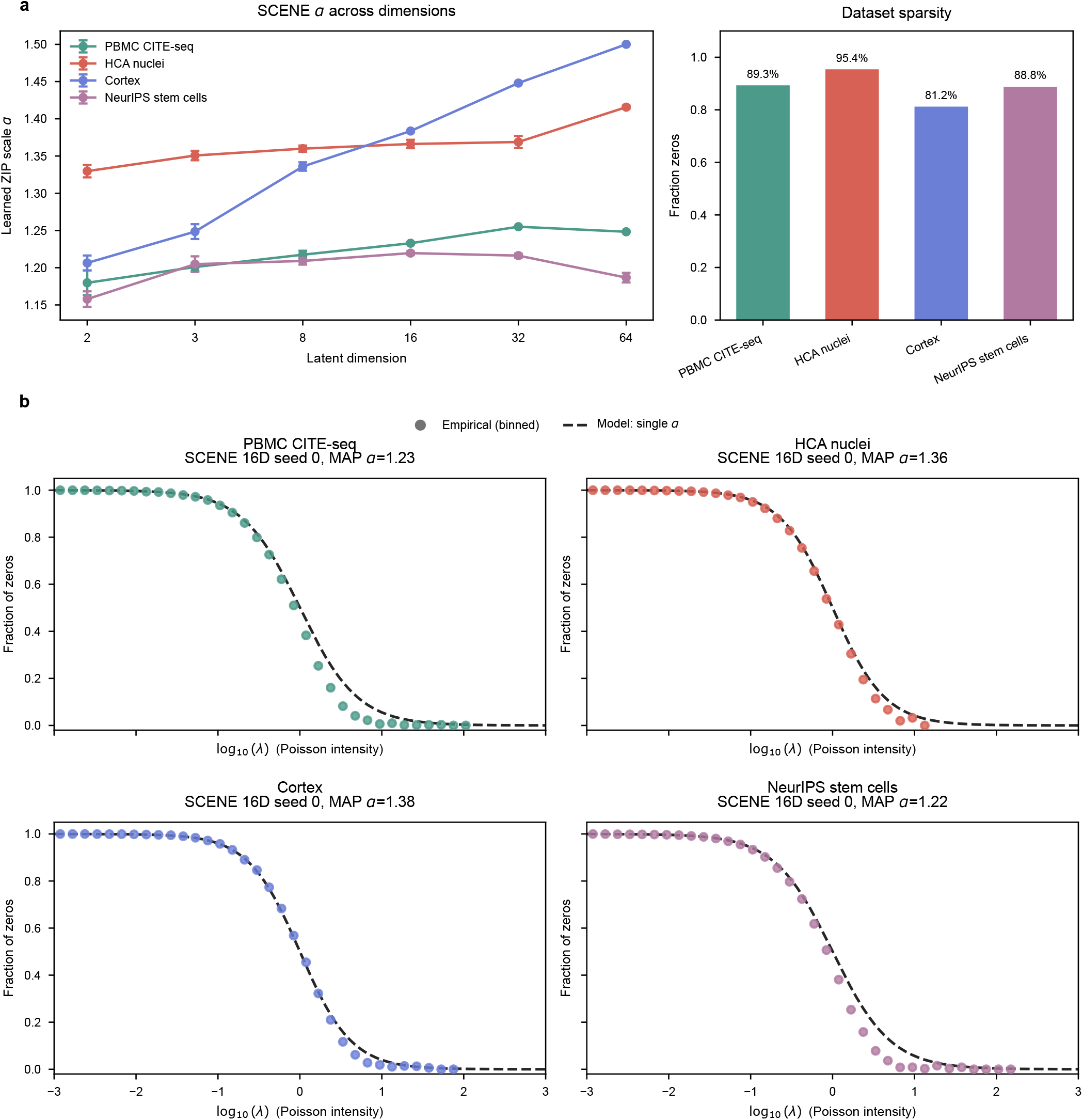
Learned zero-inflation scale and detection-intensity calibration. **a**, Learned ZIP scale parameter *α* across latent dimensionalities for each dataset, shown as mean *±* standard deviation across random seeds. The accompanying bar plot shows the empirical sparsity of each dataset as the fraction of zero entries in the count matrix. **b**, Zero fraction as a function of fitted Poisson intensity for representative 16-dimensional SCENE-ZIP models. Points show empirical zero fractions after binning cell-gene pairs by log_10_(*λ*), and dashed curves show the model-implied zero probability, 1 *− π*_*ij*_ = 1 *− σ*(*αη*_*ij*_), where *η*_*ij*_ = log *λ*_*ij*_. Agreement between empirical rates and fitted curves supports the shared monotone detection-intensity parameterization used by SCENE.

### Ultra-low-dimensional comparison

### Zero-inflation diagnostics

SCENE couples the expected positive-count intensity and the probability of nonzero expression through a shared latent predictor, with log *λ*_*ij*_ = *η*_*ij*_ and logit(*π*_*ij*_) = *αη*_*ij*_. We therefore evaluated both held-out count recovery and calibration of the learned detection-intensity relationship. Supplementary Figures 3 and 4 summarize these diagnostics across count distributions, fitted zero probabilities, and learned values of *α*.

## Supplementary Notes

### GR module Supplementary Table

**Supplementary Table 3.** Module definitions used for the GR stimulation analysis in Fig. 6a.

| Module name | Genes | Module type | Curation rationale | Expected behavior |
| --- | --- | --- | --- | --- |
| GR Direct GRE | FKBP5, SGK1, DUSP1, TSC22D3, KLF9<sup>a</sup>, PER1<sup>a</sup>, ZFP36 | Canonical glucocorticoid-responsive target genes | Compact set of established dexamethasone/GR-responsive genes selected to represent the early transcriptional response to GR activation. The module includes genes commonly used as readouts of GR pathway activation, including stress-response, phosphatase, circadian, RNA-stability, and transcriptional-regulatory targets. | Expected to show earlier contraction in gene space after dexamethasone exposure. |
| GR Late TF | KLF9<sup>a</sup>, KLF15, CEBPD, PER1<sup>a</sup>, NR4A1, NR4A2, TFAP4 | GR-associated transcription factors/regulators | Compact set of transcription factors and transcriptional regulators implicated downstream of GR activation. The module was selected to capture later regulatory organization in the GR response. | Expected to show delayed contraction in gene space compared to the GR Direct GRE module. |
| Glycolysis | HK1, GPI, PFKP, ALDOA, GAPDH, PGK1, ENO1, PKM | Canonical metabolic reference module | Compact core glycolysis module selected from central glycolytic enzymes. Included as a pathway-level metabolic reference rather than a GR-specific response module. | Expected to show compactness largely independently of the GR perturbation response; included as a stable reference program across the time course. |
| TCA cycle | CS, ACO2, IDH3A, OGDH, SUCLG1, SDHA, MDH2 | Canonical metabolic reference module | Compact core tricarboxylic acid cycle module selected from central mitochondrial TCA-cycle enzymes. Included as a mitochondrial metabolic reference rather than a GR-specific response module. | Expected to show compactness largely independently of the GR perturbation response; included as a stable reference program across the time course. |

<sup>a</sup> KLF9 and PER1 were retained in both GR modules because they are canonical glucocorticoid-responsive regulators with roles spanning direct GR target-gene induction and downstream transcriptional-response organization. Their inclusion in both modules reflects this biological overlap in GR-response architecture.

### Curation of gene modules used for the GR stimulation analysis

For the glucocorticoid receptor (GR) stimulation analysis in Fig. 6, we defined four compact gene modules before comparing SCENE and SIMBA. The purpose of these modules was not to perform de novo pathway discovery, but to test whether predefined biological programs were organized in gene space in a manner consistent with their expected biological dynamics. Specifically, we asked whether internal gene–gene distances within each module changed in accordance with known GR-response timing across the dexamethasone time course [1, 2].

The two GR modules were curated to represent two expected temporal layers of the dexamethasone response. The GR Direct GRE module contains canonical glucocorticoid-responsive genes selected to represent early target-gene induction after GR activation, including FKBP5, SGK1, DUSP1, TSC22D3, KLF9, PER1, and ZFP36. These genes were selected as a compact set of established GR-responsive readouts rather than as an exhaustive set of all GR-regulated genes [3, 4, 5].

The GR Late TF module contains GR-associated transcription factors and transcriptional regulators selected to represent downstream regulatory organization during the later response. This module was intended to capture transcriptional-regulatory structure emerging after GR activation, rather than to assert that all included genes are direct GRE-bound GR targets. The labels “GR Direct GRE” and “GR Late TF” therefore indicate the expected relative timing and biological role of the modules, not strict exclusivity of timing or mechanism for every gene [6, 7].

KLF9 and PER1 were included in both GR modules because both genes are glucocorticoid-responsive and relevant across early and later phases of the response. They are induced during the early GR transcriptional response, while their regulatory functions support their inclusion among genes contributing to downstream transcriptional-response organization [8, 4, 5].

The glycolysis and TCA-cycle modules were included as compact canonical metabolic reference modules. Unlike the GR modules, these modules represent core metabolic pathways required for basal cellular metabolism and were therefore included to test whether pathway-level gene organization remained compact across the GR stimulation time course, largely independently of the dexamethasone perturbation response [9, 10].

### All results - TF comparison in Fig. 6d

Supplementary files SIMBA TCR.xlsx, SCENE TCR.xlsx

**Supplementary Table 4.** Run-parameter specification for model configurations underlying the empirical figures and supplementary results. Figure groups are split into subrows for panel-specific configurations; comparison panels use one row per model. The batch column records only whether batch information was used; SCENE batch keys and effect parameterizations are listed in Table 5.

| Paper |  | Run parameters |  |  |  |  |  |
| --- | --- | --- | --- | --- | --- | --- | --- |
| Fig. | Panel | Model | Dataset | Dim. | Training | Input | Batch |
| Fig. 2 | a–d | **SCENE** | PBMC CITE-seq | 32D; 2D | ZIP; 600 ep.; lr 0.05; Laplacian init.; CPU; seed 42 | Counts | No |
| Fig. 3 | a–d | **SCENE** | IFN-β stim PBMC | 16D | ZIP; 600 ep.; lr 0.05; random init.; CPU; seed 42 | Counts | No |
| Fig. 4 | a | **SCENE** | HCA Nuclei | 16D | ZIP; 600 ep.; lr 0.05; Laplacian init.; seed 42 | Counts | Mixed; see Table 5 |
|  | b | **SCENE** | Cortex; PBMC CITE-seq; HCA Nuclei; NeurIPS; SIM1; SIM2 | 16D | ZIP; 600 ep.; lr 0.05; random init.; seeds 0–4 | Counts | Mixed; see Table 5 |
|  | b | Harmony/PCA | Same as Fig. 4b | 50D | PCA/Harmony; seeds 0–4 | Normalize; log1p; scale | Mixed; see Table 5 |
|  | b | SIMBA | Same as Fig. 4b | 50D | PyTorch-BigGraph training; seeds 0–4 | Library-size norm.; log; 5 bins; max 100 | Mixed; see Table 5 |
|  | b | scVI | Same as Fig. 4b | 10D; 30D batch | ZINB; NB with batch; 400 ep.; seeds 0–4 | Counts | Mixed; see Table 5 |
|  | c | **SCENE** | HCA Nuclei | 2D; 3D | ZIP; 600 ep.; lr 0.05; Laplacian init.; seed 42 | Counts | Yes; see Table 5 |
|  | d | **SCENE** | SIM2 | 16D | ZIP; 600 ep.; lr 0.05; random init.; seed 42 | Counts | Yes; see Table 5 |
|  | d | scVI | SIM2 | 30D | NB; 2 layers; 400 ep.; seed 42 | Counts | Yes |
| Fig. 5 | a–d | **SCENE** | NeurIPS stem cells | 2D; 3D | ZIP; 600 ep.; lr 0.05; Laplacian init.; CPU; seed 42 | Counts | Yes; see Table 5 |
|  | e | **SCENE** | NeurIPS stem cells; Cortex; HCA Nuclei; PBMC CITE-seq | 2D; 3D; 8D; 16D; 32D; 64D | ZIP; 600 ep.; lr 0.05; random init.; seeds 0–4 | Counts | Mixed; see Table 5 |
| Fig. 6 | a | **SCENE** | GR time course | 3D | ZIP/default SCENE count model; 200 ep.; lr 0.05; laplacian init.; seed 42 | Counts | No |
|  | a | SIMBA | GR time course | 50D | PyTorch-BigGraph training, seed 42 | Library-size norm.; log; 5 bins; max 100 | No |
|  | b–c | **SCENE** | GR control; GR 18 h | 2D | 200 ep.; lr 0.05; random init.; seed 42 | Counts | No |
|  | d–e | **SCENE** | TCR stimulation | 16D | 600 ep. lr 0.05; random init.; seed 42 | Counts | No |
|  | d–e | SIMBA | TCR stimulation | 50D | PyTorch-BigGraph training, seed 42 | Min cells 1; library-size norm.; log; 5 bins; max 100 | No |
| Supp. | Fig. 1 | **SCENE** | NeurIPS stem cells | 2D; 3D | ZIP; 600 ep.; lr 0.05; Laplacian init.; seed 42 | Counts | Yes; see Table 5 |
|  | Fig. 1 | Harmony/PCA | NeurIPS stem cells | 2D; 3D | PCA/Harmony | Normalize; log1p; scale | Yes |
|  | Fig. 1 | SIMBA | NeurIPS stem cells | 2D; 3D | PyTorch-BigGraph training; seed 42 | Library-size norm.; log; discretized graph | Yes |
|  | Fig. 1 | scVI | NeurIPS stem cells | 2D; 3D | Batch-aware NB; 400 ep.; saved config | Counts | Yes |
|  | Fig. 2 | **SCENE-ZIP** | PBMC CITE-seq | 16D | ZIP; 600 ep.; lr 0.025; CPU; split seed 42 | Counts; 10% nonzero counts held out | No |
|  | Fig. 2 | scVI | PBMC CITE-seq | 10D | ZINB; 1 layer; 400 ep.; CUDA; split seed 42 | Counts; 10% nonzero counts held out | No |
|  | Fig. 3 | **SCENE-ZIP** | PBMC CITE-seq | 16D | ZIP; 600 ep.; lr 0.025; CPU; split seed 42 | Counts; same held-out split | No |
|  | Fig. 3 | **SCENE-Poisson** | PBMC CITE-seq | 16D | Poisson; 600 ep.; lr 0.025; CPU; split seed 42 | Counts; same held-out split | No |
| | Fig. 4 | **SCENE** | PBMC CITE-seq; HCA Nuclei; Cortex; NeurIPS stem cells | 2D; 3D; 8D; 16D; 32D; 64D | ZIP; learned $\alpha$; 600 ep.; lr 0.05; seeds 0–4; 16D seed 0 for calibration panels | Counts | Mixed; see Table 5 |

**Supplementary Table 5.**
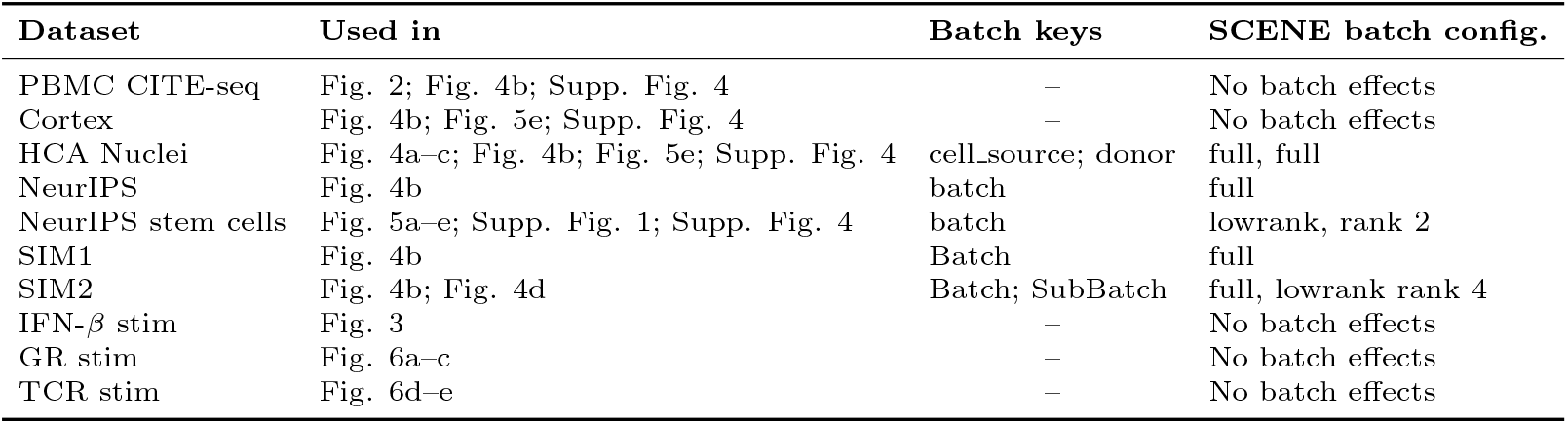
SCENE batch-effect configurations used in the empirical and supplementary analyses. For rows with multiple batch keys, entries in the configuration column follow the same order as the batch keys.

| Dataset | Used in | Batch keys | SCENE batch config. |
| --- | --- | --- | --- |
| PBMC CITE-seq | Fig. 2; Fig. 4b; Supp. Fig. 4 | – | No batch effects |
| Cortex | Fig. 4b; Fig. 5e; Supp. Fig. 4 | – | No batch effects |
| HCA Nuclei | Fig. 4a–c; Fig. 4b; Fig. 5e; Supp. Fig. 4 | cell_source; donor | full, full |
| NeurIPS | Fig. 4b | batch | full |
| NeurIPS stem cells | Fig. 5a–e; Supp. Fig. 1; Supp. Fig. 4 | batch | lowrank, rank 2 |
| SIM1 | Fig. 4b | Batch | full |
| SIM2 | Fig. 4b; Fig. 4d | Batch; SubBatch | full, lowrank rank 4 |
| IFN-β stim | Fig. 3 | – | No batch effects |
| GR stim | Fig. 6a–c | – | No batch effects |
| TCR stim | Fig. 6d–e | – | No batch effects |

